# Self-avoidance alone does not explain the function of Dscam1 in mushroom body axonal wiring

**DOI:** 10.1101/2022.04.14.488273

**Authors:** Haiyang Dong, Pengjuan Guo, Jian Zhang, Lili Wu, Ying Fu, Lei Li, Yanda Zhu, Yiwen Du, Jilong Shi, Shixin Zhang, Guo Li, Bingbing Xu, Lina Bian, Xiaohua Zhu, Wendong You, Feng Shi, Xiaofeng Yang, Jianhua Huang, Yongfeng Jin

## Abstract

Alternative splicing of *Drosophila Dscam1* into 38,016 isoforms provides neurons with a unique molecular code for self-recognition and self-avoidance. A canonical model suggests that homophilic binding of identical Dscam1 isoforms on sister branches of mushroom body (MB) axons supports segregation with high fidelity, even when only a single isoform is expressed. Here we generated a series of mutant flies with a single exon 4, 6, or 9 variant, encoding 1,584, 396, or 576 potential isoforms, respectively. Surprisingly, most of the mutants in the latter two groups exhibited obvious defects in the growth, branching, and segregation of MB axonal sister branches. This demonstrates that repertoires of 396 and 576 Dscam1 isoforms were not sufficient for normal patterning of axonal branches. Moreover, reducing Dscam1 levels largely reversed the defects caused by reduced isoform diversity, suggesting a functional link between Dscam1 expression levels and isoform diversity. Taken together, these results indicate that canonical self-avoidance alone does not explain the function of Dscam1 in MB axonal wiring.

## INTRODUCTION

Neural circuits rely on numerous cell surface receptors for target recognition, including the *Drosophila* Down syndrome cell adhesion molecule (*Dscam1*) and vertebrate clustered protocadherins (*PCDH*s).^1^ The *Drosophila melanogaster Dscam1* gene can generate 38,016 closely related single-transmembrane proteins of the immunoglobulin superfamily through mutually exclusive alternative splicing of 12 exon 4s, 48 exon 6s, 33 exon 9s, and 2 exon 17s. These comprise 19,008 ectodomains linked to one of two alternative transmembrane segments.^2^ Each ectodomain preferentially exhibits isoform-specific homophilic binding: each isoform binds only to itself, but does not bind, or only weakly binds, to other isoforms *in vivo* or *in vitro*.^3,4^ Each neuron only produces 14–50 Dscam1 isoforms through probabilistic splicing, thereby providing each neuron with a unique molecular code for self-recognition within the nervous system.^5-8^

The striking multiplicity of Dscam1 isoforms and their recognition specificity suggest that they play a unique role in the assembly of neural circuits. Genetic studies have revealed that Dscam1 is required for the wiring of diverse neurons, including mushroom body (MB) axons, dendrites from dendritic arborization (da) neurons, and olfactory projection neurons.^9-14^ A canonical model suggests that isoform-specific homophilic binding of identical Dscam1 receptor isoforms on sister branches initiates neurite repulsion, referred to as self-avoidance.^7^ Self-avoidance ensures correct spacing between sister neurites. This molecular model of self-avoidance has also been proposed for the mammalian clustered PCDHs in several neuronal populations.^15-20^

In this model, a single Dscam1 isoform is sufficient to ensure the detachment of sister self-dendrites from the same neuron.^21^ Likewise, single-cell analysis showed that a single Dscam1 isoform was sufficient to promote branch segregation with high fidelity in MB axons.^11,22^ However, to ensure that different neurons have specific identities, thousands of Dscam1 isoforms are needed to discriminate between self and non-self neurites.^23^ Although the specific isoforms expressed by a single neuron are not important, it is crucial that the subset of isoforms expressed by each neuron are different from those of its neighbors.

However, whether this canonical model provides a mechanistic basis for understanding the functions of Dscam1 isoforms in neural circuit assembly remains unclear.^24-28^ Some studies report that Dscam1 isoform diversity is also required for complex axonal branching of mechanosensory neurons.^24^ Another study investigating Dscam-RNAi knockdown in *Drosophila* motoneurons showed that Dscam1 played no role in dendrite spacing, but was essential for dendritic growth.^26^ Our recent study revealed that Dscam1 isoform bias may play a role in MB axonal wiring by influencing non-repulsive signaling.^29^ These observations suggest that Dscam1 did not function in the dendrite or axon patterning of some neurons via a self-avoidance mechanism.

Moreover, most functional experiments used a Dscam1 loss of function mutation, RNAi knockdown, or overexpression of a specific isoform.^9-13,26,30,31^ A few experiments have investigated the role of Dscam1 ectodomain diversity within the nervous system, ^11,23,25,32,33^ which mainly harbors deletions of relatively small exon 4 and 9 clusters. However, few studies have examined the effect of the number of exon 6 variants on neuronal wiring, except for mutants with a single exon 6 variant in mechanosensory axons.^24^ Therefore, whether phenotypic defects in *Dscam1* mutants are a functional requirement for specific exon clusters or isoforms, a *Dscam1* loss-of-function, or both remains unclear. Moreover, whether the diversity of exon clusters 4, 6, and 9, or their individual isoforms, is equally required for neuronal wiring remains a crucial and unresolved issue. A definitive way to determine the importance of diversity in exons 4, 6, and 9 would be to reduce the exon variants to a single exon expressed from the endogenous locus. A series of mutants expressing a single exon variant will allow comparison between isoforms and identification of differences among variable exons.

In this study, CRISPR/Cas9 technology was used to construct a series of mutant flies with a single exon 4, 6, or 9 variant, encoding 1,584, 396, or 576 potential isoforms, respectively. In contrast to previous studies,^11,23^ the mutants in the latter two groups exhibited two additional high-frequency phenotypes: truncated MB lobes and thinning of two lobes. Surprisingly, our analysis of single neurons indicated that expression repertoires of 396 and 576 Dscam1 isoforms were not sufficient for normal axonal patterning. This suggested that canonical self-avoidance alone does not explain the functions of Dscam1 in axonal wiring. These data extend the canonical model of MB in which neuron branching is based on Dscam1-mediated self-avoidance.^7,11,22,23^ Moreover, reducing Dscam1 levels largely reversed the defects caused by reduced isoform diversity, suggesting a functional link between Dscam1 expression levels and isoform diversity. These findings not only expand current understanding regarding how Dscam1 isoforms operate as cell adhesion molecules via self-avoidance, but also provide a model for studying neurological disorders involving dysregulated Dscam expression.

## RESULTS

### Mutants with a single exon 4, 6, or 9 variant

To explore how the diversity and specificity of variable exon 4, 6, and 9 clusters are differentially required for nervous system development, we constructed a series of mutants with a single exon 4, 6, or 9 (Figure 1A). First, CRISPR/Cas9 technology was combined with homologous recombination to replace the genomic region encoding variable exon 4s with complementary DNA encoding a single exon 4 (Figure S1A). The resulting 12 mutants encoded 1,584 (1 × 48 × 33) potential isoforms, referred to as *Dscam*^Single4.x^ (e.g., *Dscam*^Single4.1^) according to the remaining exon 4 variants. All mutants with a single exon 4 were homozygous viable, and fertile. Reverse transcription polymerase chain reaction (RT–PCR) using primers for exons 3 and 5 of *Dscam1* showed that two *Dscam*^Single4.5^ and *Dscam*^Single4.8^ mutants exhibited considerable abnormal splicing, due to the use of a cryptic splice site (Figure S1D). Since exon skipping affected the overall mRNA and protein level of Dscam1, phenotype experiments on these two mutants were not continued. These *Dscam*^Single4.x^ mutants lacking all but one of the exon 4 variants allowed comparison of functional differences among individual variable exons.

**Figure 1.**
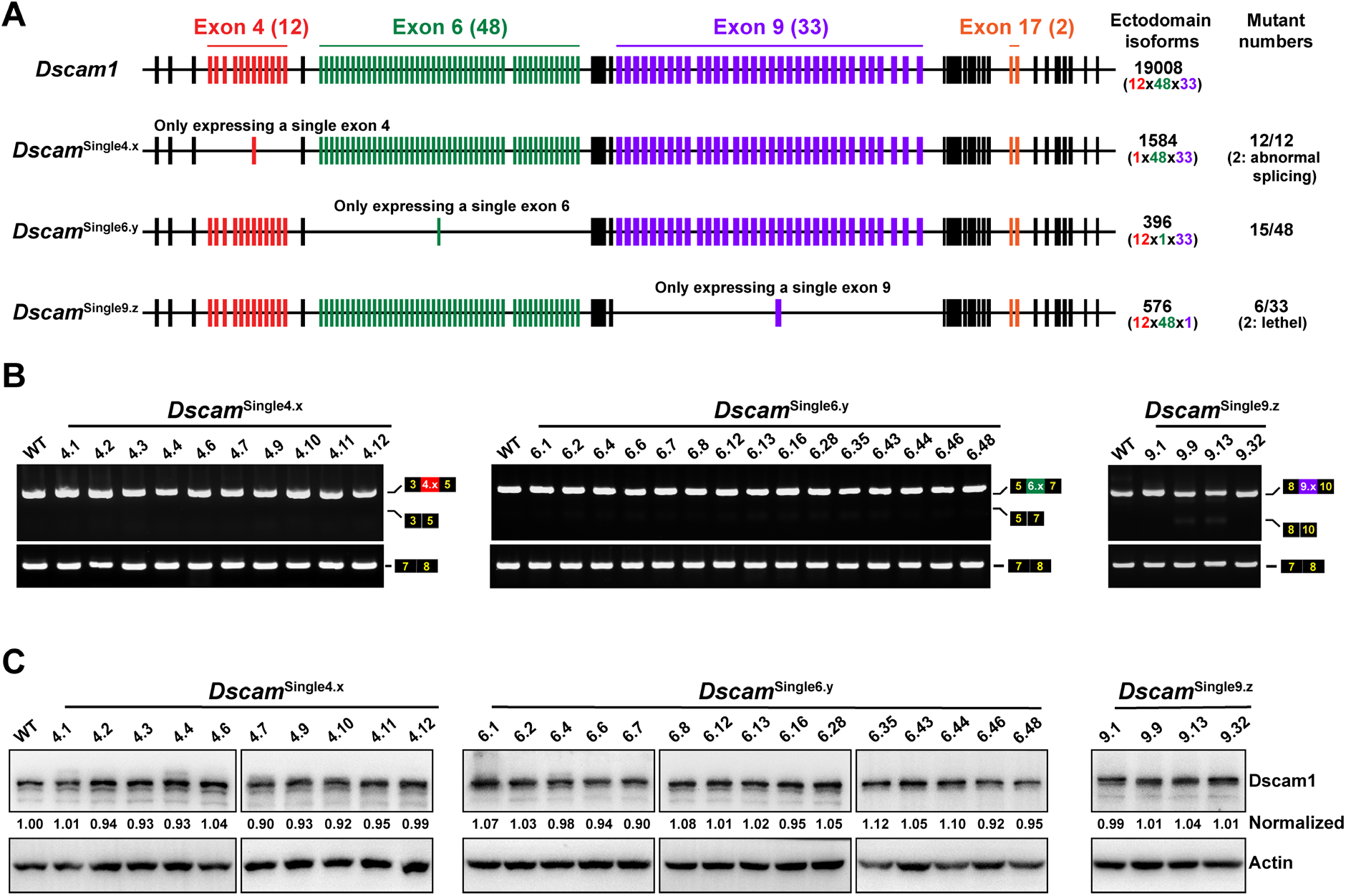
Molecular characterization of *Dscam1* deletion mutants (see also Figure S1). **(A)** Schematic diagram of mutants with a single exon 4, 6, or 9 variant. Potential ectodomain diversity; the number of mutants is shown on the right. **(B)** RT-PCR diagrams of head tissue from *Dscam1* mutants. The overall inclusion of variable exons in the mutants for each variable cluster was indistinguishable from that of the wild-type controls. **(C)** The *Dscam1* mutants had a protein level similar to that of the wild-type controls.

Mutants with a single exon 6 variant (*Dscam*^Single6.y^) that encoded 396 (12 × 1 × 33) potential isoforms were also generated using a similar method (Figure 1A; Figure S1B). Fifteen distinct exon 6s were arbitrarily selected. The selected *Dscam*^Single6.y^ mutants were homozygous viable, and fertile. RT–PCR using primers for exon 5 and 7 of *Dscam1* showed no detectable band on exon 6 skipping (Figure 1B). Several attempts have been made to generate mutants with a single exon 9 variant using a similar strategy. However, no constructs have been successfully knocked into the *Dscam1* locus. Therefore, the whole variable exon 9 cluster was replaced with genomic DNA encoding a single exon 9 (*Dscam*^Single9.z^) in this study (Figure S1C). The resulting *Dscam*^Single9.z^ mutants encoded 576 (12 × 48 × 1) potential isoforms (Figure 1A). Six distinct exon 9s (exon 9 mutants) were selected, two of which (exons 9.8 and 9.9) were the same as the mutants previously constructed by homologous recombination.^23^ In contrast to exon 4 and 6 clusters, not all six *Dscam*^Single9.z^ alleles with a single exon 9 were homozygous viable, and fertile. Homozygous viable, fertile, mutants with a single exon 9.8 and single exon 9.31 could not be generated, suggesting that *Dscam*^Single9.8^ and *Dscam*^Single9.31^ alleles were lethal recessive. RT–PCR analyses of four viable homozygous *Dscam*^Single9.z^ mutants demonstrated that exon 9 skipping was negligible (Figure 1B). Western blot assay showed that the level of Dscam1 protein in these mutants was indistinguishable from that of the wild-type (Figure 1C). These series of *Dscam*^Single4.x^, *Dscam*^Single6.y^, and *Dscam*^Single9.z^ mutants allowed study of the phenotypic consequences of reducing Dscam1 diversity and identified potential cluster- and exon variant-specific functions in parallel.

### Reducing Dscam1 diversity affects fly viability and locomotion in a cluster- and exon variant-specific manner

The effect of the diversity and specificity of variable exon 4, 6, and 9 clusters on fly viability was tested first. A mild to modest reduction in survival rate in various *Dscam*^Single4.x^ mutants encoding 1,584 isoforms was observed; the largest reduction was seen in *Dscam*^Single4.1^ mutants (Figure 2A, panel i). In contrast, all *Dscam*^Single6.y^ mutants with a single exon 6 had an obvious reduction in survival rate. Only 2–55% of embryos from *Dscam*^Single6.y^ mutants survived to adulthood (Figure 2A, panel ii), indicating that reducing Dscam1 diversity to 396 severely undermined fly viability. In addition, hatching rates in *Dscam*^Single6.y^ mutants were significantly lower compared to controls, whereas pupation and eclosion rates were not significantly affected (Figure 2A, panel ii). Moreover, the survival rate in different *Dscam*^Single6.y^ mutants varied strikingly, with the highest survival rate seen in *Dscam*^Single6.13^ flies and the lowest in *Dscam*^Single6.6^ flies.

**Figure 2.**
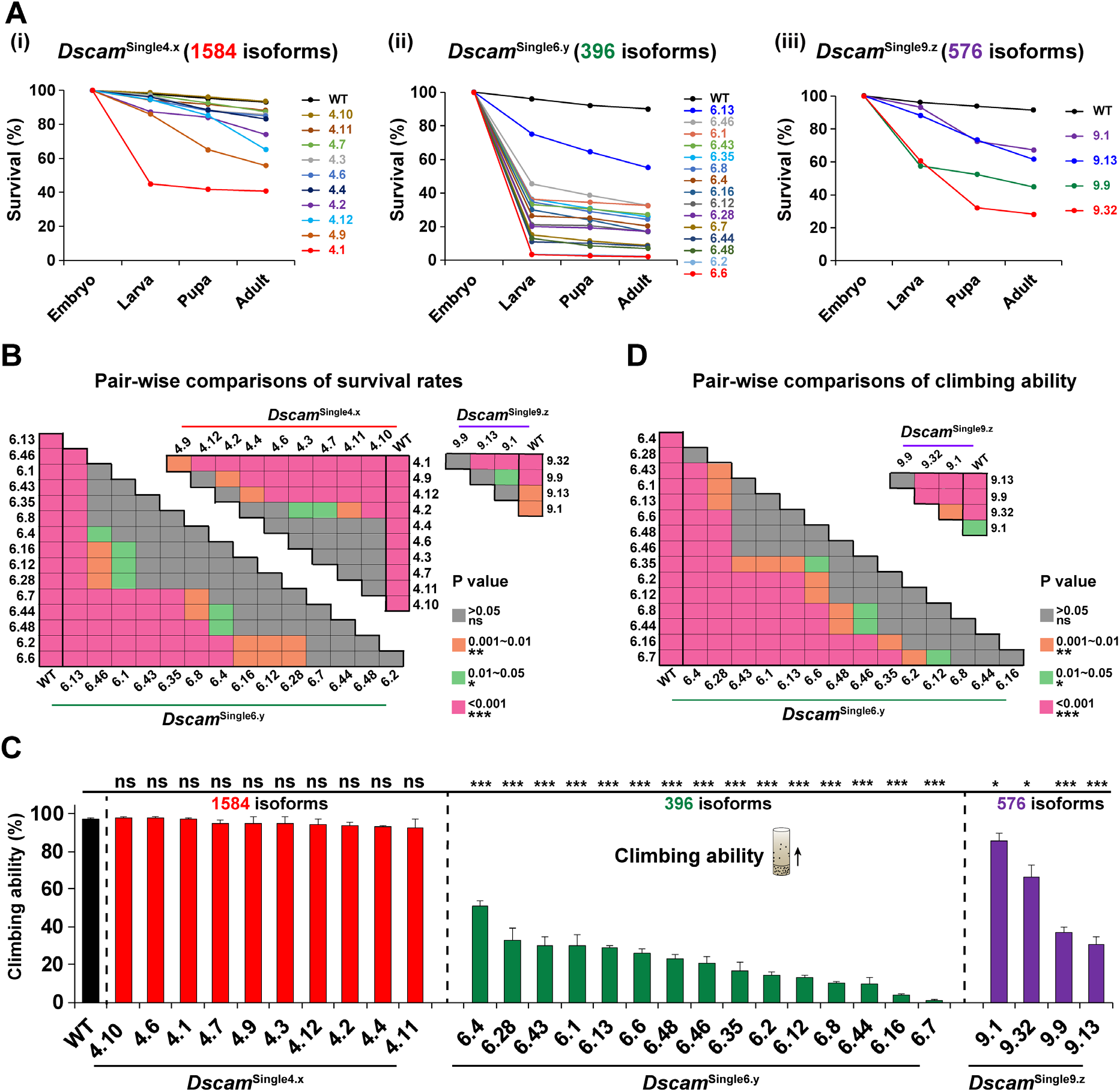
Reducing Dscam1 diversity affects fly viability and locomotion in a cluster- and exon variant-specific manner (see also Figure S2). **(A)** Survival rates of wild-type and *Dscam1* mutants during development. Survival rates at each development stage for *Dscam1* mutants expressing a single exon 4 (panel i), exon 6 (panel ii), or exon 9 (panel iii) variant. **(B)** Pair-wise comparisons (One-way ANOVA with Tukey’s test) of survival rates among *Dscam*^Single4.x^, *Dscam*^Single6.y^, or *Dscam*^Single9.z^ mutants. **(C)** Reducing Dscam1 diversity affects fly locomotion in a cluster- and exon variant-specific manner. n.s., not significant; *P < 0.05; **P < 0.01; ***P < 0.001. (Student’s t-test, two-tailed). **(D)** Pair-wise comparisons of fly locomotion among *Dscam*^Single6.y^ or *Dscam*^Single9.z^ mutants.

A similar reduction in survival rate was observed when using *Dscam*^Single9.z^ mutant alleles that encoded 576 isoforms (Figure 2A, panel iii). The survival rate in flies that carried two different *Dscam*^Single6.y^ alleles (transheterozygous animals encoding 792 isoforms) or two different *Dscam*^Single9.z^ alleles (transheterozygous animals encoding 1,152 isoforms) improved remarkably (Figure S2A). Overall, the survival rates improved as Dscam1 isoform diversity increased, but varied considerably (Figure S2B). Pairwise comparisons of the survival rates of *Dscam*^Single4.x^, *Dscam*^Single6.y^, and *Dscam*^Single9.z^ mutants indicated a discrepancy among mutants with the same isoform diversity (Figure 2B). Since *Dscam*^Single4.x^, *Dscam*^Single6.y^, and *Dscam*^Single9.z^ mutants encoded an identical number of isoforms with similar overall expression, we speculated that the discrepancy in survival rate could be likely attributed to differences in the intrinsic features of individual Dscam1 isoforms.

To assess the impact of reducing Dscam1 diversity on adult behavior, we performed a climbing assay of the mutant adults to investigate differences in locomotor abilities. No significant reduction in locomotion rate in *Dscam*^Single4.x^ mutants encoding 1,584 isoforms was observed (Figure 2C). In contrast, *Dscam*^Single6.y^ flies encoding 396 isoforms were inactive. The climbing ability of *Dscam*^Single6.y^ flies significantly decreased, by 50–98%, compared to 2-day-old control flies. The climbing ability of *Dscam*^Single9.z^ mutants encoding 576 isoforms was significantly lower than that of wild-type controls but higher than that of *Dscam*^Single6.y^ flies. The reduced climbing ability improved remarkably in flies that carried two different *Dscam*^Single6.y^ alleles encoding 792 isoforms and two different *Dscam*^Single9.z^ alleles encoding 1,152 isoforms (Figure S2C). Pairwise comparisons revealed differences in climbing ability among the *Dscam*^Single4.x^, *Dscam*^Single6.y^, and *Dscam*^Single9.z^ mutants (Figure 2D), which could be a consequence of differences in the intrinsic features of the individual isoforms. Overall, these data indicated that Dscam1 diversity was correlated with fly climbing ability (Figure S2D), suggesting that exon 6 or 9 diversity is required for normal fly locomotion in a diversity-related and variable exon-specific manner.

### Mutants with the same degree of diversity show differences in phenotypic defects in dendritic self-/non-self-discrimination of da neurons

To assess the contribution of diversity and specificity of variable exon 4, exon 6, and exon 9 clusters in nervous system development, we first assessed how the diversity and specificity of variable exon 4, exon 6, and exon 9 clusters affected overlapping of dendrites of class I and III da neurons. Previous studies have indicated that the expression of a single Dscam1 isoform was sufficient to support dendrite self-avoidance, and that expression of a common isoform among different classes of da neurons impaired overlapping of dendritic fields.^9,10,21,23^ We found that self-dendrites of class I neurons over the larval body wall seldom overlap in all *Dscam*^Single4.x^, *Dscam*^Single6.y^, and *Dscam*^Single9.z^ mutant flies (Figure S3A-C). Consistent with previous studies,^9,10,21,23^ these data indicated that expressing one exon 4, 6, or 9 variant is sufficient to support dendrite repulsion between self-branches from the same neuron.

However, different *Dscam*^Single4.x^, *Dscam*^Single6.y^, and *Dscam*^Single9.z^ mutants exhibited a considerable difference in overlapping between class I and III dendrites (Figure 3A-C). The *Dscam*^Single4.x^ mutants encoding 1,584 isoforms were indistinguishable from the wild-type flies (Figure 3B). However, compared to the wild-type, dendrites of class I and III neurons showed significantly fewer dendritic overlaps in all *Dscam*^Single6.y^ mutants encoding 396 potential ectodomain isoforms and *Dscam*^Single9.z^ mutants encoding 576 potential ectodomain isoforms (Figure 3A, C). The phenotypes of “dendritic coexistence” in flies that carried two different *Dscam*^Single6.y^ alleles encoding 792 isoforms improved remarkably, and similar results were obtained for flies carrying two different *Dscam*^Single9.z^ alleles encoding 1,152 isoforms (Figure 3C). To further assess phenotypic differences, we performed pairwise comparisons of class I and III overlapping among *Dscam*^Single6.y^ or/and *Dscam*^Single9.z^ mutants (Figure 3D). Notably, *Dscam*^Single6.y^ mutants encoding 396 potential ectodomain isoforms exhibited considerable differences in dendritic coexistence, as did *trans* heterozygous *Dscam*^Single6.y^ mutants encoding 792 potential ectodomains (Figure 3D). For example, *Dscam*^Single6.1^ had the highest penetrance of class I and III repulsion, while *Dscam*^Single6.16^ exhibited the highest level of dendritic coexistence (Figure 3C). A similar trend was observed in *Dscam*^Single9.z^ mutants, such that *Dscam*^Single9.32^ had significantly greater dendritic coexistence than the other three mutants (Figure 3C). These data imply that dendritic coexistence may be associated with intrinsic properties of individual Dscam1 isoforms. Overall, these observations indicated that Dscam1 diversity was largely correlated with class I and III dendritic coexistence (Figure 3B, C; Figure S3D). However, different mutants with the same degree of diversity showed differences in phenotypic defects associated with dendritic self-/non-self-discrimination of da neurons (Figure 3D), suggesting cluster- and exon variant-specificity.

**Figure 3.**
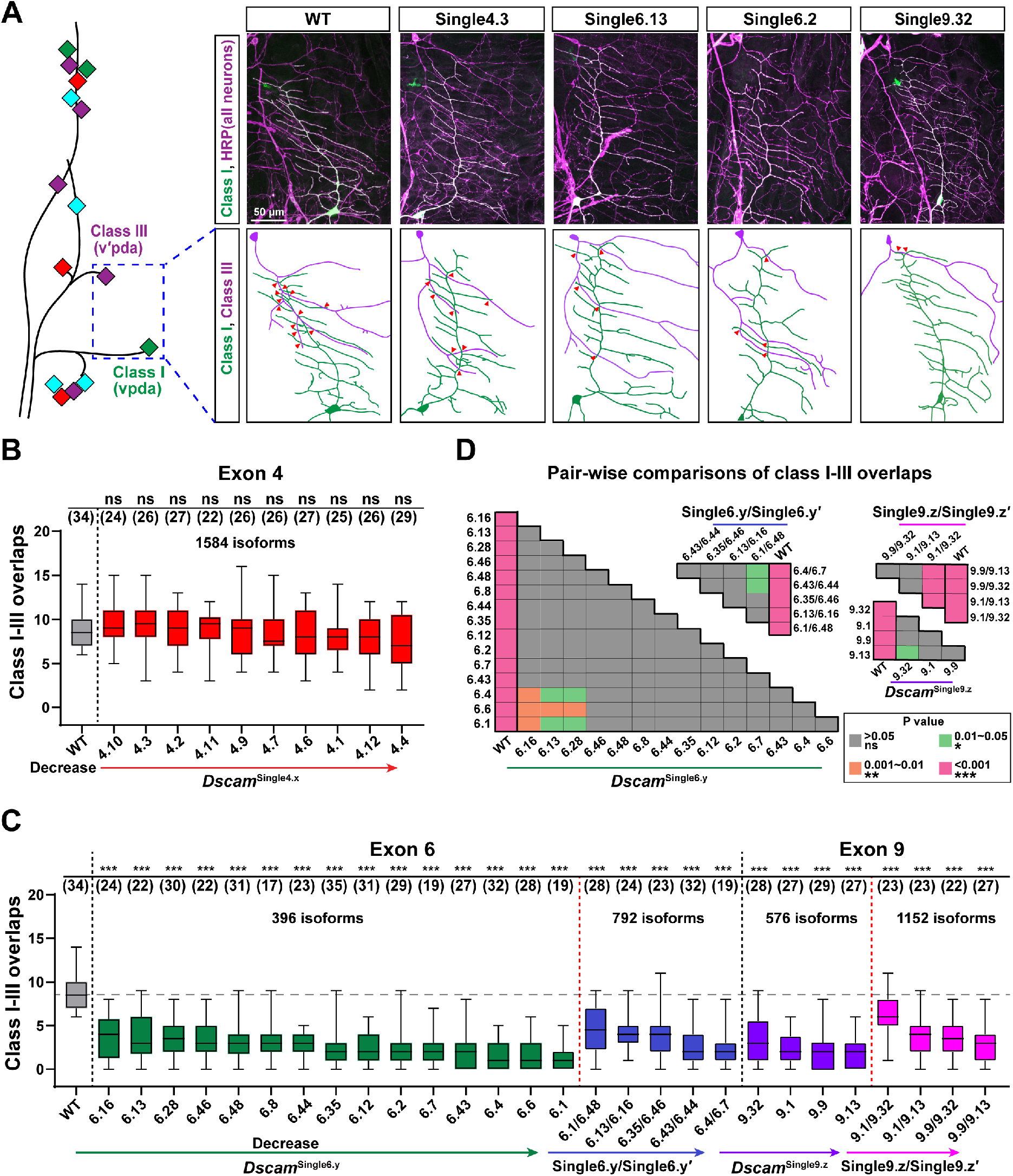
Mutants with the same degree of isoform diversity show phenotypic differences in dendritic coexistence (see also Figure S3). **(A)** Representative images of dendrites of dendritic arborization (da) neurons in wild-type and *Dscam1* mutants. A schematic diagram of the distribution of four types of da neurons (colored diamonds) is shown on the left. All neurons were visualized with an anti-HRP antibody (magenta), and class I (vpda) neurons were labeled with GFP (green; they appear white because they overlap the magenta staining). Red arrowheads indicate overlapping between class I and III dendrites. Scale bars, 50 μm. **(B)** Quantification of overlapping between class I and III dendrites in *Dscam*^Single4.x^ mutants. The numbers in parentheses represent the analyzed class I and III neurons from each genotype. **(C)** Quantification of the overlapping between class I and III dendrites in *Dscam*^Single6.y^ and *Dscam*^Single9.z^ mutants. Transheterozygous mutants, generated by combining two different *Dscam*^Single6.y^ or *Dscam*^Single9.z^ alleles, had remarkably improved phenotypes in terms of dendritic coexistence. **(D)** Pair-wise comparisons (One-way ANOVA with Tukey’s test) of overlapping class I and III dendrites among *Dscam*^Single4.x^, *Dscam*^Single6.y^, or *Dscam*^Single9.z^ mutants. These data indicate that mutants with the same degree of isoform diversity exhibited phenotypic differences in dendritic self-/non-self discrimination.

### *Dscam*^Single6.y^ and *Dscam*^Single9.z^ mutants exhibit considerably truncated MB lobes

We next assessed how the diversity and specificity of variable exon 4, 6, and 9 clusters affected the MB phenotype. Previous studies have indicated that thousands of Dscam1 isoforms are required to form normal MBs.^23^ Fas II immunostaining of normal fly brains showed that wild-type MBs consist of two orthogonal lobes of similar width that project to the dorsal lobe and towards the midline lobe, while mutants exhibited different degrees of MB phenotype defects (Figure 4A, B; Figures S4 and S5). MB phenotype defects in different *Dscam*^Single4.x^ mutants encoding 1,584 isoforms occurred at a frequency of 0–8% (Figure 4B, panel i). In contrast, *Dscam*^Single6.y^ mutants encoding 396 isoforms and *Dscam*^Single9.z^ mutants encoding 576 isoforms exhibited significant MB defects at high penetrance (64–100%) (Figure 4B, panel ii), consistent with substantially lower Dscam1 isoform diversity compared to *Dscam*^Single4.x^ mutants. Interestingly, MB phenotypes of all *Dscam*^Single6.y^*/Dscam*^*+*^ and *Dscam*^Single9.z^*/Dscam*^*+*^ heterozygous hybridization mutants were indistinguishable from those of wild-type controls (Figure S4C). The frequencies of normal MBs were significantly improved in *Dscam*^Single6.y^ transheterozygous mutants encoding 792 isoforms and *Dscam*^Single9.z^ transheterozygous mutants encoding 1,152 isoforms (Figure 4B, panel ii). Overall, our genetic data indicated that Dscam1 diversity was inversely correlated with the frequency and severity of MB phenotypic defects (Figure 4B; Figure S4D). These data support the notion that Dscam1 isoform diversity is required for normal MB development.

**Figure 4.**
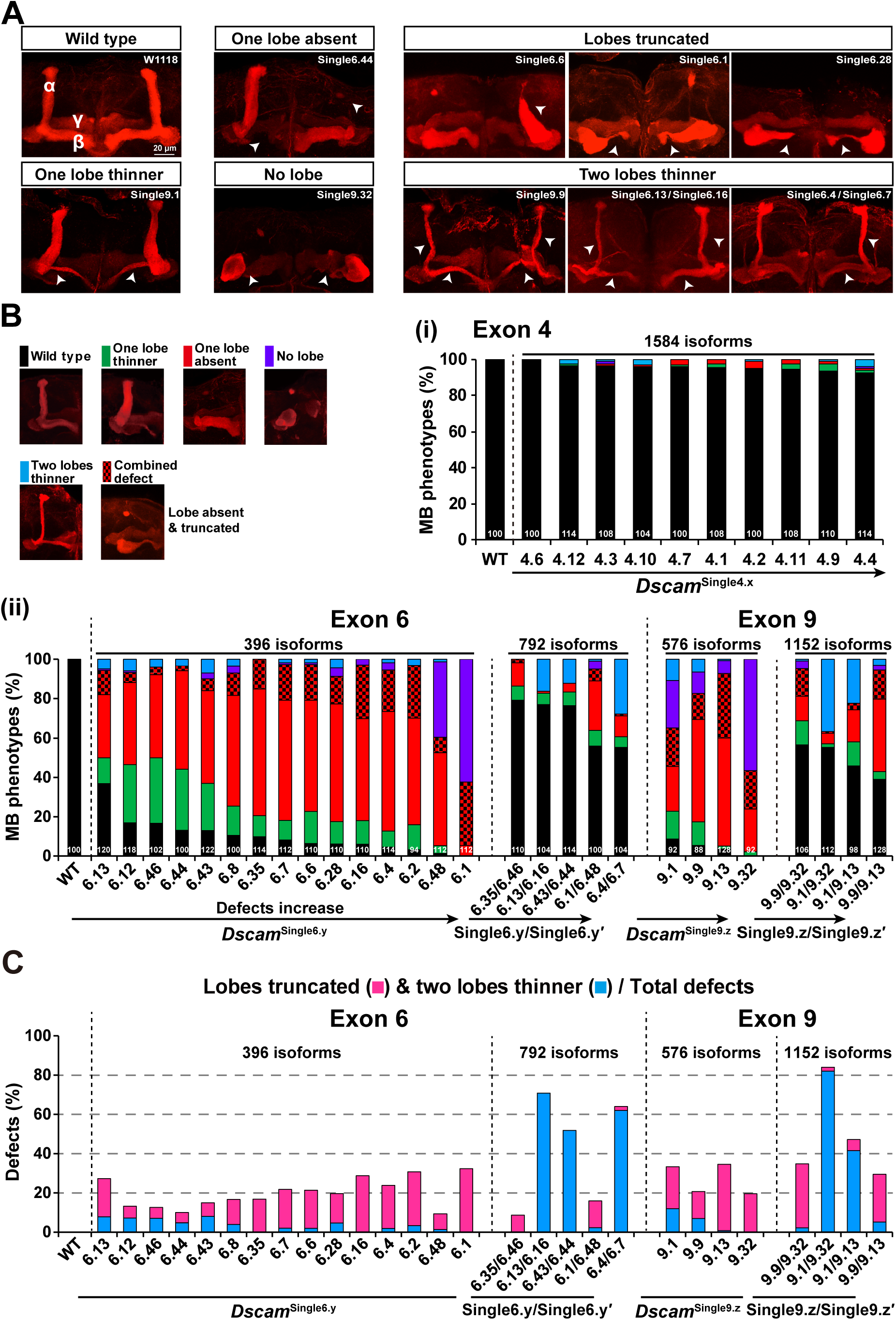
Mutants with the same degree of isoform diversity exhibit considerably different spectra of mushroom body phenotypic defects (see also Figures S4 and S5). **(A)** Mushroom body (MB) lobe morphology in wild-type and *Dscam1* mutants was visualized using monoclonal antibody 1D4 (anti-Fas II, red). See Figure S4 for other images. In addition to three defective phenotypes (no lobes, one lobe missing, one lobe thinner),^11,23^ the *Dscam*^Single6.y^ and *Dscam*^Single9.z^ mutant flies exhibited two additional phenotypes with high frequency: truncated MB lobes and thinning of two lobes. The white arrowheads indicate the defective lobes. Scale bar, 20 μm. **(B)** Quantification of MB defects in *Dscam*^Single4.x^ (panel i), *Dscam*^Single6.y^, and *Dscam*^Single9.z^ mutants (panel ii). Combining two different *Dscam*^Single6.y^ or *Dscam*^Single9.z^ alleles remarkably improved the phenotypes in terms of MBs. Numbers in parentheses refer to the numbers of MB neurons analyzed for each genotype. **(C)** Quantification of MB with lobe truncation and thinning of two lobes. The extensive lobe truncation and thinning of two lobes cannot be explained by the canonical self-avoidance model.

Notably, in contrast to previously reported *Dscam*^single-isoform^ and *Dscam*^576-isoform^ mutant flies, which only exhibited three defect types of two absent lobes, one absent lobe, or one thinner lobe,^11,23^ *Dscam*^Single6.y^ mutants exhibited a diverse range of lobe defects (Figure 4A–C; Figure S4A, B). In addition to the absent and thinned lobes, truncated MB lobes were observed in most *Dscam*^Single6.y^ mutants. This accounted for 6–32% of all defects (Figure 4B, C; Figure S4A). Likewise, MB lobes in *Dscam*^Single9.z^ mutants showing different degrees of truncation were observed (Figure 4B, C; Figure S4A). Truncated lobes were the dominant abnormalities in *Dscam*^Single6.1^ and *Dscam*^Single9.13^ mutants, which accounted for more than 30% of total defects (Figure 4C). Moreover, such lobe truncation mainly occurred in the β lobes of MBs (Figure S5A, B). Interestingly, we also observed a considerable number of mutants with thinning in two lobes. In particular, thinning in two lobes occurred extensively in transheterozygous mutants (Figure 4C; Figure S4B). For example, this accounted for more than 80% of all defects in *Dscam*^Single9.1/ Single9.32^ mutants that potentially encoded 1,152 isoforms. These two previously unreported defective MB phenotypes (lobe truncation and thinning of two lobes) accounted for 10–48% of the defects in most *Dscam*^Single6.y^ and *Dscam*^Single9.z^ mutants (Figure 4C). This cannot be explained by the canonical self-avoidance model.^11^ Since additional high-frequency phenotypes have been observed in mutants harboring different isoforms or exon clusters (Figure 4C; Figure S5A, B), such phenotypic defects should be not attributed to CRISPR-induced off-target mutations.

### Mutants with the same degree of diversity show considerably different spectra of MB phenotypic defects

While we showed that reducing Dscam1 diversity led to defective MB phenotypes, the penetrance of different mutants with the same degree of diversity varied considerably. Different *Dscam*^Single4.x^ mutants encoding 1,584 isoforms exhibited from almost no to very minor MB phenotypic defects (Figure 4B, panel i). However, *Dscam*^Single6.y^ mutants exhibited a severe yet different spectra of MB phenotype defects overall. *Dscam*^Single6.1^ had the highest penetrance and nearly all MBs exhibited obvious defects, while *Dscam*^Single6.13^ exhibited the lowest penetrance; only 64% of MBs showed defects (Figure 4B, panel ii). Similarly, different *Dscam*^Single9.z^ mutants exhibited a variety of severe phenotypic defects, and different *Dscam*^792-isoform^ transheterozygous mutants varied strikingly in MB phenotype defects, as did the various *Dscam*^1152-isoform^ transheterozygous mutants carrying two different *Dscam*^Single9.z^ alleles (Figure 4B, panel ii). Pairwise comparisons of MB defects among *Dscam*^Single6.y^ and *Dscam*^Single9.z^ mutants indicated that even mutants with the same degree of diversity exhibited considerably different spectra of MB phenotypic defects (Figure S5C). This was likely the consequence of differences in the intrinsic features of individual Dscam1 isoforms. These data also suggest that Dscam1 isoforms function to some extent in a cluster- and exon-variant-specific manner. In addition, MB phenotypes in individual *Dscam*^Singele6.y^ mutants showed a weak and non-significant correlation with class I and III dendrite overlapping (r = 0.5223, p = 0.0553; Figure S5D), suggesting that Dscam1 isoforms act on MB phenotypes via a pathway distinct from class I and III dendrite repulsion.

### *Dscam*^Single6.y^ and *Dscam*^Single9.z^ mutants exhibit axonal growth and branching defects in single-cell clones

As described above, our data showed that *Dscam*^Single6.y^ and *Dscam*^Single9.z^ animals exhibited frequent truncation of MB lobes and the thinning of two lobes. To determine the cellular phenotypes correlated with the different MB phenotypes, MARCM analysis was used to assess the axonal phenotypes at a single-cell resolution.^34^ Since *Dscam*^Single4.x^ mutants displayed only mild MB defects, we only conducted single-cell phenotypic analyses of *Dscam*^Single6.y^ and *Dscam*^Single9.z^ mutants. Unlike wild-type control clones, 41–56% of *Dscam*^Single6.y^ and *Dscam*^Single9.z^ mutant neurons exhibited either a growth defect (shorter branches than wild-type), a branching defect (single axonal branch or multiple branches), a guidance defect (e.g., axon branches deviating from the dorsal or medial direction), or a combination of the guidance/branching and growth defects (Figure 5A, B). The penetrance of the phenotypes was similar to that observed in *Dscam*^null^ flies (Figure 5B).

**Figure 5.**
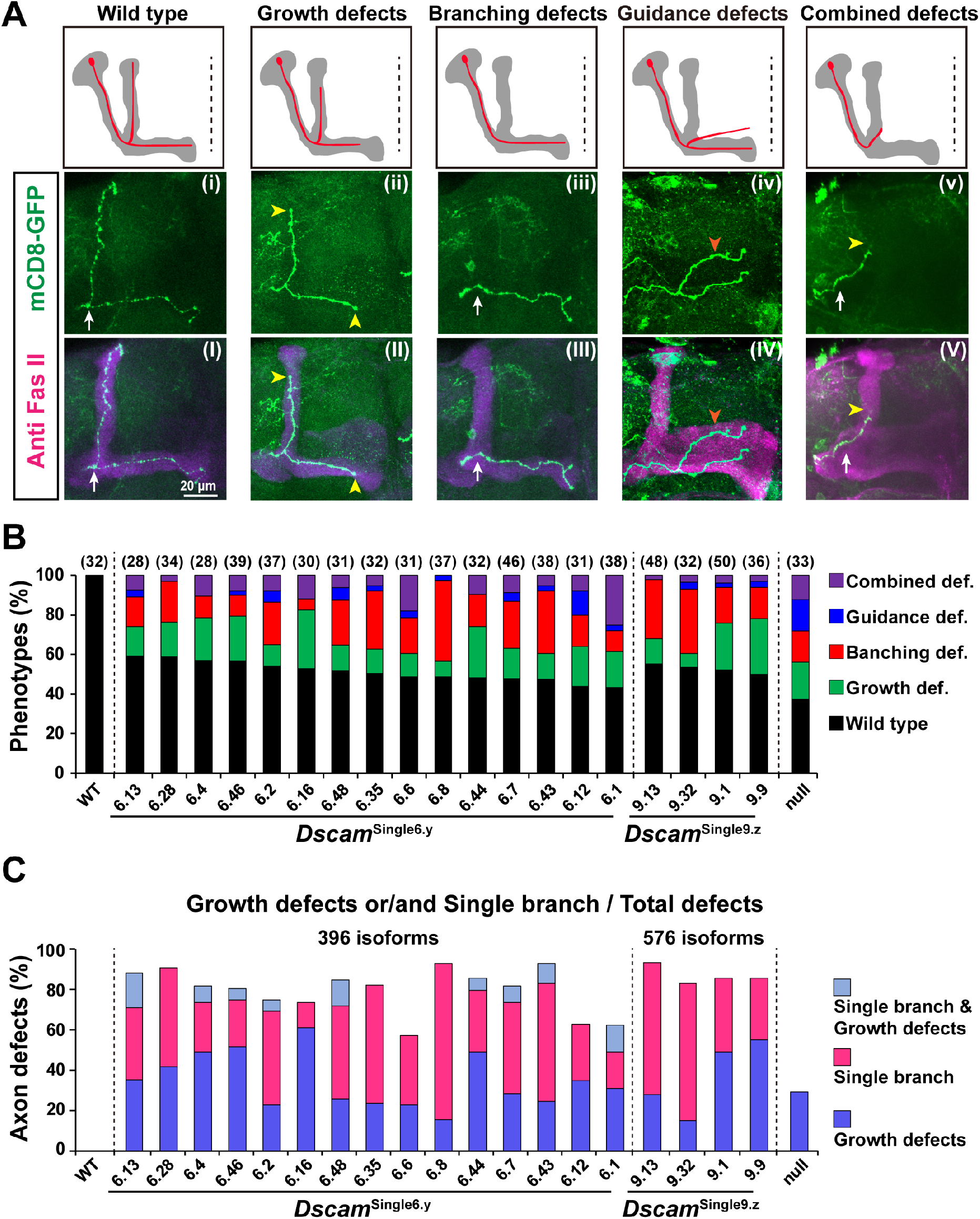
*Dscam*^*S*ingle6.y^ and *Dscam*^Single9.z^ mutants exhibit MB axonal defects in single-cell clones. **(A)** Schematic summary of different defective phenotypes in mutant MB neurons. All images are of adult brain tissue. Labeled single-cell clones were revealed by GFP (green) and MB lobes were immunostained with anti-Fas II (red). Note an unusual single-branch defect with no axon bifurcation at the peduncle end in certain single-cell clones. Scale bar, 20 μm. **(B)** Quantification of axonal defects in single-cell clones. Numbers in parentheses represent the number of single neurons in each genotype. **(C)** Growth or/and single-branch defects relative to the total number of defects. The data show that expression of a repertoire containing up to 396 or 576 Dscam1 isoforms was not sufficient for normal axonal branching and growth.

Axonal growth defects accounted for up to 15–62% of all axonal defects (Figure 5C), suggesting that axon growth was sensitive to the reduction of Dscam1 diversity. Most growth defects were in the form of shortened axon branches (Figure 5A, panel ii). This indicated that axon growth terminated prematurely or failed to reach the ends of the α and β lobes in mutant clones. The high incidence of short axons was consistent with the high frequency of truncated MB lobes observed in *Dscam*^Single6.y^ and *Dscam*^Single9.z^ mutants. Overextended axon branches (extended over the midline) were rarely observed in these mutants. These observations indicated that stunted axon growth caused extensive truncation of lobes in *Dscam*^Single6.y^ and *Dscam*^Single9.z^ mutant brains.

Axon branching showed modest sensitivity to a reduction in Dscam1 diversity, which accounted for 13–79% of all axonal defects (Figure 5C). An unusual single-branch/neuron phenotype, where α- but not β-axonal branches were generated without axon bifurcation at the peduncle end and vice versa, was detected. The single-branch/neuron phenotype varied remarkably among various single-cell MARCM mutant clones (Figure 5C). The single-branch neuron phenotype accounted for up to 70% of *Dscam*^Single6.8^ mutant single-cell clones. In contrast, only 18% of mutant axons failed to bifurcate at the peduncle terminus in *Dscam*^Single6.1^ single-cell mutant clones (Figure 5C). These observations indicated that Dscam1 diversity potently modulated axon bifurcation. Branching defects showing a single branch might result from collateral retraction due to strong homophilic interactions at the bifurcate. The absence, thinning, or thickening of one lobe in *Dscam*^Single6.y^ and *Dscam*^Single9.z^ mutants may be attributed to the suppression of axon bifurcation or misguidance of the bifurcated branches. We observed that axon bifurcation at the peduncle terminus was suppressed for some mutant single-cell clones (Figure 5A, panel iii), while two bifurcated axonal branches were commonly segregated into an α- or β- lobe for others. These findings indicated that the loss and thinning of a lobe, as observed in fly mutant whole brains (Figure 4A), was caused by a combination of axon branching failure and misguidance.

We also observed axon misguidance in *Dscam*^Single6.y^ and *Dscam*^Single9.z^ single-cell mutant clones. Specifically, two bifurcated axonal branches failed to segregate reliably into α- or β- lobes (Figure 5A, panel iv). This may have been due to incomplete retraction as a result of moderately strong homophilic interactions. Indeed, most of the guidance defects appeared in combination with growth defects. Taken together, these data showed that even repertoires containing 396 and 576 Dscam1 isoforms were not sufficient to sustain proper axonal patterning.

### *Dscam*^Single6.y^ and *Dscam*^Single9.z^ mutants do not exhibit reliable segregation in MB axonal branches

A fundamentally different role for Dscam1 diversity in regulating MB branch segregation was proposed in previous studies,^11,22,31^ where axonal sister branch segregation of MB neurons was analyzed in different *Dscam*^single^ mutants encoding a single ectodomain. In their study, the sister branches of isolated *Dscam*^single^ MB neurons segregated with high fidelity, whereas the sister branches of *Dscam*^null^ neurons did not. These findings led to the proposal that, consistent with dendritic patterning, the expression of single isoforms was sufficient to restore sister branch segregation at the single-cell level, such that Dscam1 diversity is dispensable within a single neuron. Here, the role of Dscam1 diversity in sister branch segregation of MB neurons is revisited. In contrast to previous work,^11,22,31^ our data demonstrated that even potential repertoires of 396 and 576 Dscam1 isoforms (encoded within the genome of a single cell) were not sufficient for reliable segregation into α and β branches.

In contrast to the nearly normal segregation pattern seen in wild-type axons, bifurcated sister branches in *Dscam*^Single6.y^ mutant neurons may run in diverse directions in single-cell clones (Figure 6A). Interestingly, bifurcated sister branches in a few neurons segregated normally at the beginning and extended in the dorsal or medial direction, but later, their extension deviated from the normal trajectory. Statistical analyses showed that the phenotypes varied remarkably between different *Dscam*^Single6.y^ mutants encoding 396 isoforms (Figure 6B). Moreover, the frequency of segregation defects in most *Dscam*^Single6.y^ mutants was not significantly different from that of *Dscam*^null^ mutants. Notably, some *Dscam*^Single6.y^ mutants exhibited significantly greater segregation defects compared with wild-type controls (Figure 6B). For example, > 20% of *Dscam*^Single6.1^ axons with bifurcated sister branches were not reliably segregated into the dorsal and medial lobes, as observed in the *Dscam*^null^ mutants (Figure 6B). In addition, we found no significant difference of branch segregation defects between *Dscam*^null^ and *Dscam*^Single9.z^ mutants (Figure 6B). Similar phenomena have been reported regarding the axonal branching of mechanosensory neurons.^24^ Taken together, our data revealed that expressing a repertoire of 396 or 576 Dscam1 isoforms was not sufficient for branch segregation with high fidelity at the single-cell level.

**Figure 6.**
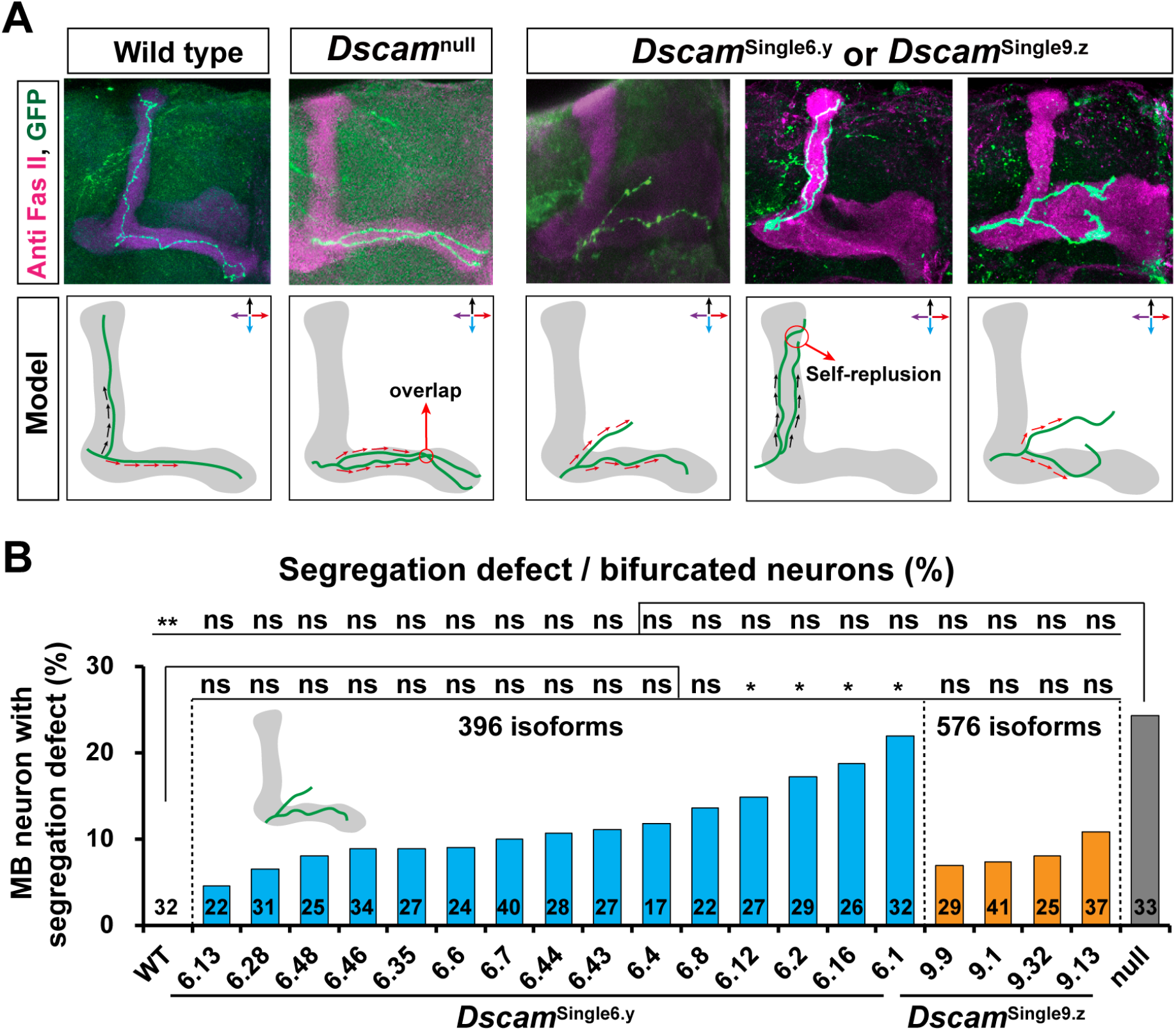
*Dscam*^Single6.y^ and *Dscam*^Single9.z^ exhibit segregation defects of MB axonal branches. **(A)** Schematic showing diverse segregation defects of MB axonal branches in *Dscam*^Single6.y^ and *Dscam*^Single9.z^ mutants. Arrows indicate the extension direction of axonal branches. **(B)** Quantification of MB neurons with abnormal branch segregation in single-cell clones. These data show that the frequencies of segregation defects in *Dscam*^Single6.y^ and *Dscam*^Single9.z^ mutants were indistinguishable from those of the *Dscam*^null^ mutant. *P < 0.05; **P < 0.01; ***P < 0.001; n.s., not significant (Fisher’s exact test, two-tailed).

### Lowering Dscam1 levels in *Dscam*^Single6.y^ and *Dscam*^Single9.z^ mutants largely reverses defects caused by reduced isoform diversity

To further elucidate the molecular mechanisms underlying defective phenotypes, we investigated the effect of Dscam1 levels on developmental and neuronal defects in *Dscam*^Single6.y^ and *Dscam*^Single9.z^ flies. We compared *Dscam*^Single6.y^/*Dscam*^null^ and *Dscam*^Single9.z^/*Dscam*^null^ flies bearing one copy of a mutant allele, which reduced the Dscam1 expression level by approximately 50% to wild-types (Figure S6A). Analysis of fly viability revealed that as many as 40–79% of the *Dscam*^Single6.y^/*Dscam*^null^ flies survived to adulthood, compared to 2–55% of the *Dscam*^Single6.y^ homozygous flies (Figure 7A; Figure S6B). Similar trend has been observed in *Dscam*^Single9.z^ mutants. Moreover, lowering Dscam1 levels in both *Dscam*^Single6.y^ and *Dscam*^Single9.z^ mutants helped restore climbing ability (Figure 7A; Figure S6B). These observations indicated that reducing Dscam1 levels largely reversed fly viability and locomotion defects caused by reduced isoform diversity.

**Figure 7.**
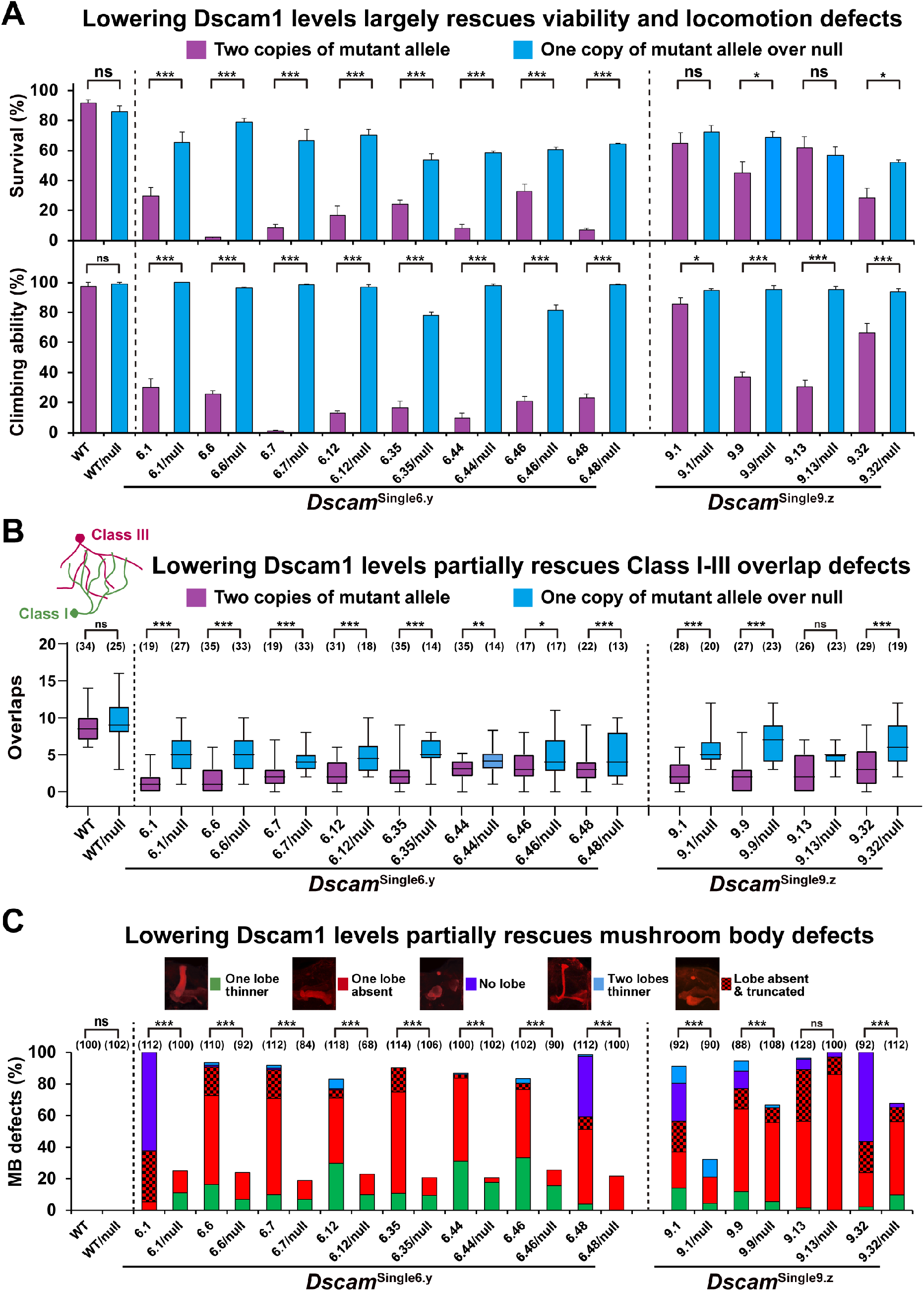
Reducing Dscam1 expression levels in *Dscam*^Single6.y^ and *Dscam*^Single9.z^ mutants partially rescues defects caused by reduced diversity (see also Figures S6 and S7). **(A)** Reducing the Dscam1 expression level partially rescued the diminished fly viability (upper panel) and climbing ability (lower panel) in homozygous mutants. **(B)** Reducing the Dscam1 expression level remarkably improved dendritic coexistence in homozygous mutants. The numbers in parentheses represent the number of mutants investigated. **(C)** Reducing the Dscam1 expression level rescued defects caused by reduced diversity of homozygous mutants. The numbers in parentheses represent the numbers of mutants investigated. Data are expressed as mean ± SD. *P < 0.05; **P < 0.01; ***P < 0.001; n.s., not significant (Student’s t-test, two-tailed).

We next investigated the effect of reducing Dscam1 levels on neuronal phenotype. We found that self-dendrites of class I neurons seldom overlap in *Dscam*^Single6.y^/*Dscam*^null^ and *Dscam*^Single9.z^/*Dscam*^null^ mutants, as was the case for *Dscam*^Single6.y^ and *Dscam*^Single9.z^ mutants (Figure S6C). However, dendrites of class I and III neurons overlapped much more in *Dscam*^Single6.y^/*Dscam*^null^ mutants than in homozygous mutants with two copies of *Dscam*^Single6.y^ (Figure 7B; Figure S7A, B). Similarly, dendrites from class I and III neurons in *Dscam*^Single9.z^/*Dscam*^null^ flies overlapped considerably compared to the minimal overlap seen in *Dscam*^Single9.z^ homozygous flies, although this was not the case for *Dscam*^Single9.13^ flies (Figure 7B). However, we found a small and statistically significant increase in dendrite overlapping of class I and III neurons in *Dscam*^Single9.13^/*Dscam*^null^ flies compared with *Dscam*^Single9.13^ homozygous flies. The self-binding affinity of the exon 9.13-encoding Ig7 domain was 2–4 times higher than those of the other three Ig7 domains.^4^ Thus, reducing the Dscam1 expression level by 50% was still not sufficient to significantly reverse defective dendritic overlapping. Taken together, our results indicated that repulsion behavior between dendritic neurites is not only determined by the number of Dscam1 isoforms, but is also associated with Dscam1 protein levels and self-binding affinity.

Moreover, the normal MB phenotype rate in *Dscam*^Single6.y^/*Dscam*^null^ mutants improved by up to 80%, whereas almost no normal MB phenotypes were seen in *Dscam*^Single6.y^ flies (Figure 7C; Figure S7C). Similarly, lowering the Dscam1 levels substantially reduced the frequency of defective MBs in *Dscam*^Single9.y^ mutants, except *Dscam*^Single9.13^ mutants (Figure 7C). The latter exception may be due to the much higher self-binding affinity of the Dscam1 isoform encoding exon 9.13.^4^ However, we observed lower severity of abnormal lobe phenotypes in *Dscam*^Single9.13^/*Dscam*^null^ compared to *Dscam*^Single9.13^ mutants, although the overall frequency of defective MBs was indistinguishable between the groups (Figure 7C). Considering that the rescue effects were conserved in exon 6 and 9 clusters among different phenotypes, and also in different isoforms, these results indicated a general functional link between Dscam1 expression levels and isoform diversity.

## DISCUSSION

This study provided conclusive evidence that self-avoidance alone does not explain the functions of Dscam1 in MB axonal wiring. A CRISPR/Cas9 system was used to construct a series of mutant flies with a single exon 4, 6, or 9 variant. Analysis of single neurons indicated that a repertoire of 396 or 576 Dscam1 isoforms was not sufficient for normal patterning of axonal branches. Moreover, reducing Dscam1 levels in mutants rescued the developmental and neuronal defects caused by reduced isoform diversity. These findings expand the current understanding of how Dscam1 isoforms operate as a cell adhesion molecule via self-avoidance, and have implications for neurological disorders associated with dysregulated Dscam expression.

### Self-avoidance alone does not explain the functions of Dscam1 in MB axonal wiring

The findings in this study extend the canonical model of Dscam1-mediated self-avoidance underlying the axonal branching of MB neurons.^1,7,11,22,23^ In this model, sister branches recognize each other through Dscam1 isoform-specific homophilic binding, thereby activating a repulsion signal to segregate axons into separate pathways. However, the genetic evidence presented herein extend this mechanistic model.

First, in addition to the previous observation that all three *Dscam*^single^/*Dscam*^null^ mutants carry a single ectodomain and three *Dscam*^576-isoform^ mutant animals exhibit three types of defective phenotype (no lobes, one missing lobe, one thinner lobe), ^11,23^ the *Dscam*^Single6.y^ and *Dscam*^Single9.z^ mutant flies in this study exhibited two additional high frequency phenotypes: truncated MB lobes and thinning of two lobes (Figure 4A, B). Similarly, other mutant flies with reduced Dscam1 diversity (e.g., *Dscam*^Single6.4^/*Dscam*^Single6.7^) frequently exhibited thinning of two lobes (Figure 4B, C). These newly observed phenotypic defects could not be explained by Dscam1-mediated self-avoidance. The discrepancies may be attributable to the different mutants examined. Notably, our genetic analyses showed that the severity (phenotype) of MB defects was inversely correlated with Dscam1 diversity, and that severity was also affected by the overall Dscam1 level and properties of the individual variable Ig domain, such as self-binding affinity. Thus, compared with previous work,^11,23^ the series of mutants with different degrees of Dscam1 diversity and Dscam1 levels in this study more reliably reflected the possible phenotypes associated with MB defects.

Second, mosaic analysis revealed frequent axonal growth and branching defects in *Dscam*^Single6.y^ and *Dscam*^Single9.z^ flies. In particular, unusual single-branch phenotypes were detected in 13–79% of *Dscam*^Single6.y^ and *Dscam*^Single9.z^ mutants (Figure 5B, C). These results are reminiscent of observations in some single-cell clones rescued by overexpression of specific Dscam1 isoforms, where axon bifurcation at the peduncle terminus was suppressed in 3–13% of mutant single-cell clones.^35^ However, *Dscam*^null^ mutants do not seem to exhibit this type of single branch defect, instead exhibiting supernumerary branches at the peduncle terminus or other positions.^36^ Thus, in contrast to the loss of function seen in *Dscam*^null^ mutants, the axon bifurcation defect in *Dscam*^Single6.y^ and *Dscam*^Single9.z^ mutants might be caused by gain-of-function effects due to a marked increase of certain isoforms in MB neurons. In *Dscam*^Single6.y^ and *Dscam*^Single9.z^ mutants, reduced isoform diversity might cause excessive homophilic binding between self-branches at the initially bifurcated growth cones. The over-strong homophilic interaction likely caused one branch retraction, followed by bifurcation suppression at the peduncle terminus. Alternatively, this process may involve other functions of a single isoform, as indicated by the considerable differences in defect frequency among *Dscam*^Single6.y^ and *Dscam*^Single9.z^ mutants. The results reported here support and extend a previously proposed model that explained the coordination of axon bifurcation with divergent segregation according to Dscam1-dependent growth cone collapse.^36^ Further comparisons of MB axon growth cone among wild-type and mutant flies will be needed before it is possible to confirm the role of Dscam1 diversity in axon bifurcation.

Moreover, the axon-shortening defects observed in *Dscam*^Single6.y^ and *Dscam*^Single9.z^ mutants could not be explained by Dscam1-mediated self-avoidance. Combined with genetic evidence, we propose that Dscam1 diversity mediates MB axonal growth largely through its influence on non-repulsive signaling. Excessive signaling within MB axons is detrimental to axon growth and branching, which is relieved by a reduction in Dscam1 levels and/or increase in diversity to weaken signaling. The unusually high expression levels of single isoforms in *Dscam*^Single6.y^ or *Dscam*^Single9.z^ mutants could cause excessive homophilic interactions, which may result in the retraction of one newly formed branch. Thus, Dscam1 diversity is required for MB axon branch formation, which might buffer Dscam1 signaling to control axonal branching and growth. Further studies will be required to define filopodia dynamics during MB axon growth.

Finally, the results presented herein indicated that *Dscam*^Single6.y^ and *Dscam*^Single9.z^ mutants exhibit striking differences in the degree of MB axon branch segregation defects, some of which (e.g., *Dscam*^Single6.1^) have a similar spectrum to *Dscam*^null^ mutants. Our data are somewhat inconsistent with the previously proposed notion that the expression of single isoforms is sufficient to support sister branch segregation with high fidelity at the single-cell level.^11,22,31^ This discrepancy may be attributable to the different mutants, the different Dscam1 protein level, or to the different methods in the generation of MARCM clones. In their study ^11,22,31^, using intragenic MARCM avoided the potential cross-reactivity between MARCM cells and background cells, but MARCM cells only expressed half of the Dscam1 protein, while conventional MARCM used in this study appears to cause the potential cross-reactivity. However, our recent studies indicated the isoform-composition-altered mutants, which avoid the potential cross-reactivity between MARCM cells and background cells, exhibited obvious axonal branch segregation defects of MB neurons.^29,37^ These data suggest that there is no causal relationship between cross-reactivity and axonal segregation defects. Taken together, all lines of evidence suggest that self-avoidance alone does not explain the functions of Dscam1 in normal MB axonal growth, branching, and segregation. Instead, Dscam1 isoforms can also control normal MB axonal patterning via a non-repulsive mechanism.

### Functional link between Dscam1 expression levels and isoform diversity

This study showed that reducing the Dscam1 expression level in *Dscam*^Single6.y^ and *Dscam*^Single9.z^ flies remarkably decreased the behavioral and neuronal defects caused by reduced Dscam1 diversity. Mechanistically, *Dscam*^Single6.y^ and *Dscam*^Single9.z^ mutants expressed a significantly less diverse set of isoforms, leading to excessive homophilic binding strength. Reducing overall Dscam1 expression levels would return single isoforms to sub-toxic levels. The second possible mechanism involves other potential functions of single isoforms. Several studies have indicated that Dscam receptors recognize distinct and specific ligands.^38-40^ This possibility is partially supported by our observation that reducing overall Dscam1 expression levels modulated behavioral and neuronal defects in a variant-related manner. As Dscam1 protein could initiate signaling underlying multiple functions via homophilic and heterophilic interactions,^6,27,28,41^ Dscam1 expression levels are functionally linked to isoform diversity via combinatorial mechanisms.

The resulting over-strong Dscam1 signal likely leads to developmental and neuronal defects, as reported in previous studies.^42,43^ By contrast, reducing Dscam1 levels prevents excessive Dscam1 signaling, thus rescuing developmental, behavioral, and neuronal defects. Excessive signaling is detrimental to axon growth and branching, which is relieved by the weakened signaling induced by reducing Dscam1 levels and/or increasing diversity. In this sense, Dscam1 isoform diversity produced by alternative splicing buffers Dscam1 signaling to ensure normal neuronal development. These data suggest a functional and mechanistic link between Dscam1 levels and isoform diversity independent of isoform specificity. It will be interesting to determine whether this expression level-diversity link exists in other genes.

Appropriate regulation of Dscam expression is required for normal neuronal development; dysregulated Dscam expression may cause developmental abnormalities and diseases.^42,44,45^ In flies, Dscam1 expression levels seem to serve as an instructive code that controls the size of the presynaptic arbor.^42,43,46^ Notably, Dscam expression is elevated in several neurological disorders, such as Down syndrome,^47^ bipolar disorder,^45^ and intractable epilepsy.^44^ Dscam deficiency accelerated spine maturation and promoted autism-like behaviors.^48^ A previous study showed that Dscam expression levels correlated with presynaptic arbor size, and that excessive presynaptic arbor growth caused by elevated Dscam levels may contribute to the pathogenesis of brain disorders.^42^ Reducing Dscam levels in Fragile X mutants reduced synaptic targeting errors and rescued behavioral responses.^43^ Given the strong correlation between Dscam1 expression levels and isoform diversity seen in this study, these findings provide a model for studying neurological disorders associated with dysregulated Dscam expression and screening potential drugs for Down syndrome and other similar conditions.

## MATERIALS AND METHODS

### Materials: Fly Stocks

We used *{nos-Cas9}attP2*, delivered via in embryo microinjections, to generate germline mutations.^49^ *W*^*1118*^ was used as a wild-type control. *if/CyO* and *if/CyO*.*GFP* lines were used as the balancer stocks. The *Gal4*^*221*^ line was used to drive *UAS-mCD8-GFP* expression in class I da neurons, as described previously.^50^ *FRT42D* and *hsFLP,UAS-mCD8-GFP;FRT42D,tubP- Gal80/CyO;Gal4-OK107* lines were used to label single-cell clones of MB neurons. All flies were cultured on a standard cornmeal medium at 25°C.

### Generation of fly Dscam1 mutant alleles

We used a CRISPR/Cas9-mediated homology-directed repair system to generate the desired mutants with only one exon variant.^49,51^ In this system, two sgRNA and one donor plasmid were co-injected into *{nos-Cas9}attP2* embryos. This procedure was carried out by UniHuaii Co., Ltd. (Guangdong, China). Using DNA Extraction Kit (AG21009, ACCURATE BIOTECHNOLOGY, HUNAN, Co., Ltd.) to obtained DNA of female offspring from the crosses of injected mosaic flies and balancer stock flies were and identified by PCR analysis. Then, the male offspring from the mutation tubes, identified via PCR, were crossed with the balancer stock. DNA was extracted from every male fly for PCR analysis to acquire the final mutation lines. Next, the heterozygous offspring were collected and crossed to produce homozygous offspring. To exclude any effect of genetic background and avoid off-target effects, all mutants were backcrossed with *W*^*1118*^ lines for five generations. The insertion sites were confirmed by DNA sequencing. The primers for sgRNA and mutant screening are listed in Table S1 and the mutation sequences of the *Dscam*^Single9.z^ mutants are listed in Table S2.

### RT–PCR

According to the manufacturer’s instructions, total RNA from fly heads was extracted using TRIzol reagent (Invitrogen, Waltham, MA, USA). Reverse transcription was performed using the SuperScript III system (Invitrogen) and specific primers for *Dscam1* exon 10. The reverse transcription products (cDNA) were amplified using PrimeSTAR DNA polymerase (TaKaRa, Shiga, Japan) and specific primers. The final products were analyzed by electrophoresis with 1.5% agarose gel and photographed using the GIS 1D Gel Image System (ver. 3.73; Tanon, Shanghai, China).

### Western blot analyses

Protein was obtained from fly heads using strong RIPA lysis buffer (CW-BIO, Shanghai, China) and a protease inhibitor PMSF (Beyotime, Jiangsu, China). Western blotting was performed according to the manufacturer’s protocol (Abcam, Cambridge, UK). First, protein samples from wild-type and mutant flies were separated using 12% SDS–PAGE gel and transferred onto nitrocellulose membranes. After blocking with 5% bovine serum albumin (BSA; VWR Life Science, Radnor, PA, USA) for 1 hour at room temperature, the membranes were incubated with diluted primary antibodies to Dscam1 (ab43847; diluted 1:5,000) or β-actin (ab8227; diluted 1:10,000) at 4°C overnight. After three washes with TBST, the membranes were incubated with secondary antibodies (goat anti-rabbit IgG, 1:10,000; CW-BIO) at room temperature 2 two hours. The immunoreactive bands of Dscam1 and β-actin protein were detected using the eECL Western Blot Kit (CW-BIO) and 5200 imaging system (Tanon Science & Technology Co. Ltd., Shanghai, China).

### Growth and development

About 120 virgin female and male fruit flies were collected. All flies were mixed for 2 days in a bottle containing a juice tray and yeast extract. The juice tray was replaced every 4 hours, a total of three times. We collected 200 embryos from each juice tray. After 48 hours, the hatched embryos were counted to obtain the hatching rate. We collected 90 larvae from the hatched group of second-instar larvae and distributed them into three new food tubes (three groups in total). After 3–4 days, we counted the pupae on the tube wall and calculated the pupation rate. After another 4–5 days, we counted the adult flies in the food tube and calculated the eclosion rate.

### Climbing ability

One-day-old male flies were collected and separated into three tubes (30 flies per tube). A height of 2 centimeters was marked on each tube. The tubes were tapped so that all of the flies could be gathered at the bottom. Ten seconds after tapping, the flies that had climbed over the 2- centimeter height marker were counted. This process was repeated 10 times, and the average of 10 trials for each tube was recorded as the final result. Finally, climbing ability was measured (percentage of flies that climbed over the 2-centimeter height marker in each tube).

### MARCM clones of MB neurons

Single-cell clones of MB neurons from wild-type and *Dscam1* mutants were generated using MARCM techniques.^34^ First, *Dscam*^Single6.y^ and *Dscam*^Single9.z^ mutants were crossed with FRT42D lines to permit natural chromosome recombination. Next, *Dscam*^Single6.y^ and *Dscam*^Single9.z^ mutant males with FRT42D sites were mated with *hsFLP,UAS-mCD8- GFP;FRT42D,tubP-Gal80/CyO;Gal4-OK107* lines. Then, mitotic recombination was induced by heat shock via a 37°C water bath for 1 hour during the pupation stage. Finally, straight-winged offspring were dissected and immunofluorescence stained.

### Immunostaining

The green fluorescent protein (GFP)-labelled class I da neurons from third-instar larvae were dissected to assess the dendritic co-existence of class I and class III da neurons. The larval epidermises were fixed with 4% paraformaldehyde for 25 minutes at room temperature and blocked in 5% BSA (diluted in phosphate buffered saline tween-Tween [PBST]; phosphate-buffered saline containing 0.1% Triton X-100) after three washes in PBST (20 minutes each time). Then, a horse radish peroxidase (HRP) antibody (Cy3-conjugated Affinipure Goat Anti-HRP, diluted 1:200) was used to incubate the larvae epidermises at 4°C overnight. After three washes in PBST for 20 minutes each, the samples were mounted in ProLong Gold Antifade reagent (Invitrogen). The immunofluorescence staining was imaged using a LSM800 confocal microscope (Carl Zeiss, Oberkochen, Germany). Images of class I (vpda) neurons were exported in a single channel. For MB immunofluorescence staining, brains of 3- to 5-day-old adult flies were dissected in PBS and fixed in 4% paraformaldehyde at room temperature for 35 minutes. After three 20-minute washes with PBST, the brains were blocked with 5% BSA for 1 hour at room temperature. After another three washes, the brains were incubated for 48 hours at 4°C with primary antibodies (anti-FasII, DSHB, diluted 1:2 in PBST; anti-GFP, diluted 1:600 in PBST). After standard washing, the brains were incubated for 4 hours with secondary antibodies (Alexa594-goat-anti-mouse IgG diluted 1:400 in PBST; EarthOx, Millbrae, CA, USA; Alexa 488-goat-anti-mouse IgG, diluted 1:400 in PBST; EarthOx) at room temperature. The brains were then washed again and mounted in ProLong gold Antifade reagent. Finally, MB immunofluorescence staining was imaged using an LSM800 laser scanning confocal microscope. Images were exported using Zeiss Zen Software (version 2.3).

### Statistical analysis

Quantitative analyses were performed for three biological replicates. Error bars represent mean ±SD. Significant differences were determined using a two-tailed Student’s t-test, with p > 0.05 considered non-significant (n.s.). One-way ANOVA with Tukey’s test was used to conduct pairwise comparisons of phenotypes between different Dscam1 mutants which was performed using GraphPad Prism version 8.0.2 for Windows, GraphPad Software, San Diego, California USA. To compare with the previous data,^11^ Fisher’s exact test was used to analyze the defects of MB axons with branch segregation in single-cell clones. Pearson’s correlations between the different Dscam1 mutant phenotypes were calculated using SPSS software (version 22.0, IBM, New York, USA).

## ACKNOWLEDGMENTS

We thank Haihuai He for comments on the manuscript; and members of the Jin lab for suggestions and discussion during the course of this work. This work was supported by research grants from the National Key Research and Development Program of China (2021YFE0114900), the National Natural Science Foundation of China (91940303, 91740104), the Natural Science Foundation of Zhejiang Province (LD21C050002), and the Starry Night Science Fund at Shanghai Institute for Advanced Study of Zhejiang University (SN-ZJU-SIAS- 009).

## AUTHOR CONTRIBUTIONS

YJ conceived of this project. HD, PG, JZ, and LW designed and performed the experiments; HD, PG, JZ, LW, YF, LL, YZ, YD, JS, LB, and XZ performed generating altered bias alleles; HD, PG, JZ, LW, SH and JH performed phenotype analysis. YJ, HD, PG, JZ, BX, GL, WY, FS, XY and JH analyzed the data; YJ, HD, PG and JZ wrote the manuscript; all authors discussed the results and commented on the manuscript.

## SUPPLEMENTARY MATERIALS

Figures S1 to S7

Tables S1 to S2

