## Supplemental File for "Self-avoidance alone does not explain the function of Dscam1 in mushroom body axonal wiring"

This PDF file includes:

Tables S1 to S2

Figures S1 to S7

| Table S1 Specific primers used for sgRNA and mutant screening | | |
|---|---|---|
| **Mutants** | **Screening primers** | **sgRNA Primers** |
| ***Dscam***$^{Single4.x}$ | **Ds-4-exa-F:** TGAAATGGCTGGAACAATAGTC **Ds-4-exa-R:** CTTCAGCTATTGTGTTCTTC | **Ds-3-sg-F1 :** TTGTTATTCAAGTGCAGGGA **Ds-3-sg-R1 :** TCCCTGCACTTGAATAACAA **Ds-5-sg-F1 :** CCTGGTTGGCCAATATGCCA **Ds-5-sg-R1:** TGGCATATTGGCCAACCAGG |
| ***Dscam***$^{Single6.y}$ | **Ds-6-exa-F:** GCTACCAGTGCCGAACCAAACATC **Ds-6-exa-R:** AGTCTCAACGCTTTCGCCTCCCAC | **Ds-4.12-sg-F1 :** GGCAAGTCAGGAGCGACCCA **Ds-4.12-sg-R1 :** TGGGTCGCTCCTGACTTGCC **Ds-8-sg-F1 :** CCCAGAATCTGGGAATGCTC **Ds-8-sg-R1 :** GAGCATTCCCAGATTCTGGG |
| ***Dscam***$^{Single9.z}$ | **Ds-9-exa-F:** TATTCCCAGAGTTCCCCAGAAT **Ds-9-exa-R:** CGAGAAAAACTGGATTAGCTGC | **Ds-9.1-sg-F1 :** TCAAAGGGTACGACCTGCGG **Ds-9.1-sg-R1 :** CCGCAGGTCGTACCCTTTGA **Ds-9.33-sg-F1 :** CATTCACAGTTAGCCTGGTC **Ds-9.33-sg-R1 :** GACCAGGCTAACTGTGAATG |
| **Specific primers used for RT-PCR** | | |
| **Primers** | **5'-3' sequence** | **Assay** |
| **Ds-3-F** | TGGATCAGGAGCGACGGTAC | RT-PCR |
| **Ds-5-R** | CTCCAGAGGGCAATACCAGG | RT-PCR |
| **Ds-5-F** | GCTACCAGTGCCGAACCAAACATC | RT-PCR |
| **Ds-7-R** | AGTCTCAACGCTTTCGCCTCCCAC | RT-PCR |
| **Ds-7-F** | AACATAACCTCGGTCCACGC | RT-PCR |
| **Ds-8-R** | GTCGCTTGGTCTGAGTTCCG | RT-PCR |
| **Ds-8-F** | ACTTGCGTTGCCAAGAATCAGGAAG | RT-PCR |
| **Ds-10-R** | GCCTTATCGGTGGGCTCGAGGATCC | RT-PCR |
| **Ds-10-RT** | GGTTTGGGGAAGCCATCAGCCTT | RT |

| Table S2 Summary of the mutation sequences of *Dscam*$^{\text{Single9.z}}$ mutants | | |
|---|---|---|
| **Mutants** | **Mutation sequences*** | **Assay** |
| ***Dscam***$^{\text{Single9.1}}$ | caatcgaatacccaaacctctcgctc~~agTTCCACCGCAGGTCGTACC......**Exon9.1**......~~GTCGA<br>Ccattatgcaaatcatactcatagttaaacgtttttctattttaaattatttatttgagtttt<br>aatttaccacttatttcctgtccatttaagTTCCACCGCAGGTCGTACCCTTTGATTTCGGTG<br>AGGAAACCATCAACATGAATGACATGGTCTCGGCCACGTGCACAGTGAACAAGGGCGACACTC<br>CCCTGGAGCTGTACTGGACAACGGCTCCGGATCCCACGACGGGAGTGGACGCCTGATGTCCA<br>ACGATGGCATTCTAATCACAAAGACGACGCAGCGCATCAGCATGCTGAGCATAGAGTCCGTGC<br>ATGCTCGCCATCGGGCGAACTACACGTGTGTGGCCAGGAATGCGGCCGGGGTCATCTACCACA<br>CGGCAGAGCTGCGCGTTAACG**Exon9.1**gtttgtccccacaagaaatattgaataaggagaaa<br>taagagcacgcttactgtctatgttgttttcacttagtattgcttgaggtgtttggtgttgga<br>gcagtcgcattcatatgatatttgtgctgaatgtcatataaatcagaaaaattaggtgtaata<br>tgattattaagacaagctacaatactttaagaagaaatattctgaggagctgttttgtcattt<br>tggtagataaaatacgaccgcgttctctttgcccattcgcatgcTCTAGA~~............CAGGCTAAC<br>TGTGAATG**Exon9.33**gta~~aatgggcggttatcagattatcag | *Dscam*$^{\text{Single9.1}}$ mutant was constructed. |
| ***Dscam***$^{\text{Single9.8}}$ | caatcgaatacccaaacctctcgctc~~agTTCCACCGCAGGTCGTACC......**Exon9.1**......~~GTCGA<br>Ccattatgcaaatcatactcatagttaaacgtttttctattttaaattatttatttgagtttt<br>aatttaccacttatttcctgtccatttaagTTCCTCCCCAAATCCAAGCGTTTGACTTTGGTT<br>CTGAAGCGGCCAATACTGGTGAAATGGCGGGTGGCTTTTGCATGGTTCCCAAGGGCGATCTGC<br>CAATGGAGATCCGCTGGACCCTGAATTCAGCACCCATCATCACCGGTGAACATGGATTTTCCC<br>TGTCGCGCCTTAATCCACGTACCAGCTCCTTGAGCATCGACTCTCTCGAGGCCAGACATCGCG<br>GACTGTACCGATGTATTGCCTCCAACAAGGCGGGCAGCGCTGAATACAGCGCTGAATTGCACG<br>TGAATG**Exon9.8**gtttgtccccacaagaaatattgaataaggagaaataagagcacgcttac<br>tgtctatgttgttttcacttagtattgcttgaggtgtttggtgttggagcagtcgcattcata<br>tgatatttgtgctgaatgtcatataaatcagaaaaattaggtgtaatatgattattaagacaa<br>gctacaatactttaagaagaaatattctgaggagctgttttgtcattttggtagataaaatac<br>gaccgcgttctctttgcccattcgcatgcTCTAGA~~............CAGGCTAACTGTGAATG**Exon9.3**<br>**3**gta~~aatgggcggttatcagattatcag | *Dscam*$^{\text{Single9.8}}$ mutant was constructed. |
| ***Dscam***$^{\text{Single9.9}}$ | caatcgaatacccaaacctctcgctc~~agTTCCACCGCAGGTCGTACC......**Exon9.1**......~~GTCGA<br>Ccattatgcaaatcatactcatagttaaacgtttttctattttaaattatttatttgagtttt<br>aatttaccacttatttcctgtccatttaagTGCCGCCCCAGGTTTTGCCCTTTAGTTTCGGCG<br>AATCCGCCGCCGATGTCGGCGATATTGCCAGTGCCAACTGTGTGGTGCCCAAGGGAGATCTGC<br>CCCTGGAGATTCGCTGGTCCCTCAACTCTGCTCCAATCGTAAACGGCGAAAATGGGTTTACCC<br>TGGTGCGTCTGAATAAGCGCACGAGTCTGTTAAACATTGATTCTCTAAATGCTTTTCATCGCG<br>GTGTCTACAAGTGCATAGCCACAAATCCGGCTGGGACCAGTGAATATGTAGCTGAATTACAAG<br>TTAATG**Exon9.9**gtttgtccccacaagaaatattgaataaggagaaataagagcacgcttac<br>tgtctatgttgttttcacttagtattgcttgaggtgtttggtgttggagcagtcgcattcata<br>tgatatttgtgctgaatgtcatataaatcagaaaaattaggtgtaatatgattattaagacaa<br>gctacaatactttaagaagaaatattctgaggagctgttttgtcattttggtagataaaatac<br>gaccgcgttctctttgcccattcgcatgcTCTAGA~~............CAGGCTAACTGTGAATG**Exon9.3**<br>**3**gta~~aatgggcggttatcagattatcag | *Dscam*$^{\text{Single9.9}}$ mutant was constructed. |
| ***Dscam***$^{\text{Single9.13}}$ | caatcgaatacccaaacctctcgctc~~agTTCCACCGCAGGTCGTACC......**Exon9.1**......~~GTCGA<br>Ccattatgcaaatcatactcatagttaaacgtttttctattttaaattatttatttgagtttt<br>aatttaccacttatttcctgtccatttaagTTTTGCCCCAAATTGTGCCCTTCGCCTACGAGG<br>ATCTGATCAATATGGGCGACTCGATAGATTTGTTTTGCCAAATCCAAAAAGGCGACCGTCCCA<br>TCAAGGTGCACTGGAGTTTCGAGCGGAGCGCTGGAGACTACGGCTTTGATCAGGTGCAGCCCC<br>AGATGCGCACGAATCGCATTAGCGAGAAGACGAGCATGATCTCCATTCCCAGTGCCAGTCCTG<br>CCCACACCGGCCGGTATACCTGTATAGCCAGTAATAAGGCTGGAACTACAACATATAGCGTTG<br>ACCTGACAGTGAACG**Exon9.13**gtttgtccccacaagaaatattgaataaggagaaataaga<br>gcacgcttactgtctatgttgttttcacttagtattgcttgaggtgtttggtgttggagcagt<br>cgcattcatatgatatttgtgctgaatgtcatataaatcagaaaaattaggtgtaatatgatt<br>attaagacaagctacaatactttaagaagaaatattctgaggagctgttttgtcattttggta<br>gataaaatacgaccgcgttctctttgcccattcgcatgcTCTAGA~~............CAGGCTAACTGTGA<br>ATG**Exon9.33**gta~~aatgggcggttatcagattatcag | *Dscam*$^{\text{Single9.13}}$ mutant was constructed. |

| | | |
|---|---|---|
| ***Dscam***$^{\mathbf{Single9.31}}$ | caatcgaatacccaaacctctcgctc~~agTTCCACCGCAGGTCGTACC……**9.1**……~~GTCGACcat<br>tatgcaaatcatactcatagttaaacgtttttctattttaaattatttatttgagttttaatt<br>taccacttatttcctgtccatttaagTGCCCCCCAAACTGGCTCCCTTGCCCGTAAATTCGCC<br>ACTGTACGTGGGCGACTACTACCAGTTGACGTGCGCGGTGGTGCACGGCGACGCACCCTTCAA<br>CATTACCTGGTACTACAACAACGAGCCCGCCGGAGACTTGGCCGGGGTCACCATCCTGATGCA<br>CGGCCGGCGCAGCAGCTCGCTGAACATCGAGAGCGTGGGGGGTGACCACGCGGGCAACTACAC<br>CTGCAAGGCCGCCAACCGGGCGGGCGAAACCACGGCTGAGACCCATCTGAGTGTCAAGG**9.31**<br>gtttgtccccacaagaaatattgaataaggagaaataagagcacgcttactgtctatgttgtt<br>ttcacttagtattgcttgaggtgtttggtgttggagcagtcgcattcatatgatatttgtgct<br>gaatgtcatataaatcagaaaaattaggtgtaatatgattattaagacaagctacaatacttt<br>aagaagaaatattctgaggagctgttttgtcattttggtagataaaatacgaccgcgttctct<br>ttgcccattcgcatgcTCTAGA~~…………CAGGCTAACTGTGAATG**Exon9.33**gta~~aatgggcgg<br>ttatcagattatcag | *Dscam*$^{\text{Single9.31}}$ mutant was constructed. |
| ***Dscam***$^{\mathbf{Single9.32}}$ | caatcgaatacccaaacctctcgctc~~agTTCCACCGCAGGTCGTACC……**Exon9.1**……~~GTCGA<br>Ccattatgcaaatcatactcatagttaaacgtttttctattttaaattatttatttgagtttt<br>aatttaccacttatttcctgtccatttaagAGCTGCCCCAGGTTGATCAGTTCCACTTCAATG<br>CCAACGGGGTGAATGGCGGCCAGGCAGTGCGGGTCATGTGCATGGTCTCCTCCGGCGACCTGC<br>CCATCGACATCTACTGGCTGAAGGACGGACAGCCCCTGCTGCGCTCCATCTACCACAAGATCG<br>ACGAATACACCCTGATCCTGTCGCTGCGTCAGACCACCATTGGTGACTCCGGCAACTACACGT<br>GCGTGGCCAGCAACGCGGCGGGCGTGGCCAGTCGCTGGTCCATCCTGAAAGTGAAGG**Exon9.**<br>**32**gtttgtccccacaagaaatattgaataaggagaaataagagcacgcttactgtctatgttg<br>ttttcacttagtattgcttgaggtgtttggtgttggagcagtcgcattcatatgatatttgtg<br>ctgaatgtcatataaatcagaaaaattaggtgtaatatgattattaagacaagctacaatact<br>ttaagaagaaatattctgaggagctgttttgtcattttggtagataaaatacgaccgcgttct<br>ctttgcccattcgcatgcTCTAGA~~…………CAGGCTAACTGTGAATG**Exon9.33**gta~~aatgggc<br>ggttatcagattatcag | *Dscam*$^{\text{Single9.32}}$ mutant was constructed. |

*The variable exon sequences are in capital letters and highlighted in green. Intron sequences are lowercase letters. The sequences in red font indicate the mutation sequences. The sequences shown in red font and strikethrough represent the deleted sequences.

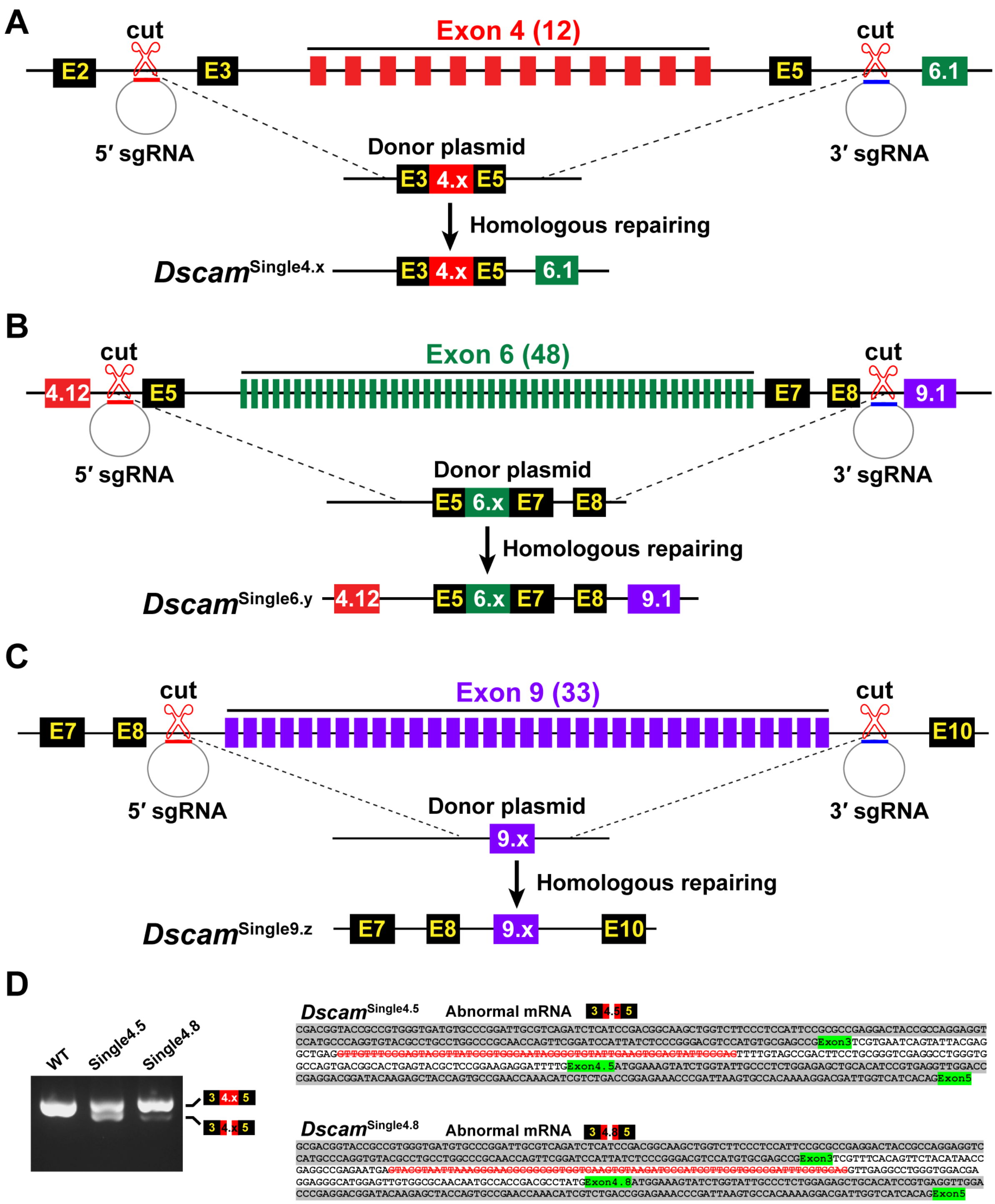


**Figure S1. Schematic diagram of the construction of *Dscam1* mutant flies. (Related to Figure 1).**

**(A-C)** Schematic diagram of the construction of mutants with a single variable exon 4, 6 and 9. **(D)** RT-PCR diagram from the heads of wild-type and *Dscam*$^{\text{Single4.x}}$ mutant flies. RT-PCR and sequencing showed that two *Dscam*$^{\text{Single4.5}}$ and *Dscam*$^{\text{Single4.8}}$ mutants exhibited considerable abnormal splicing, due to the use of a cryptic splice site.

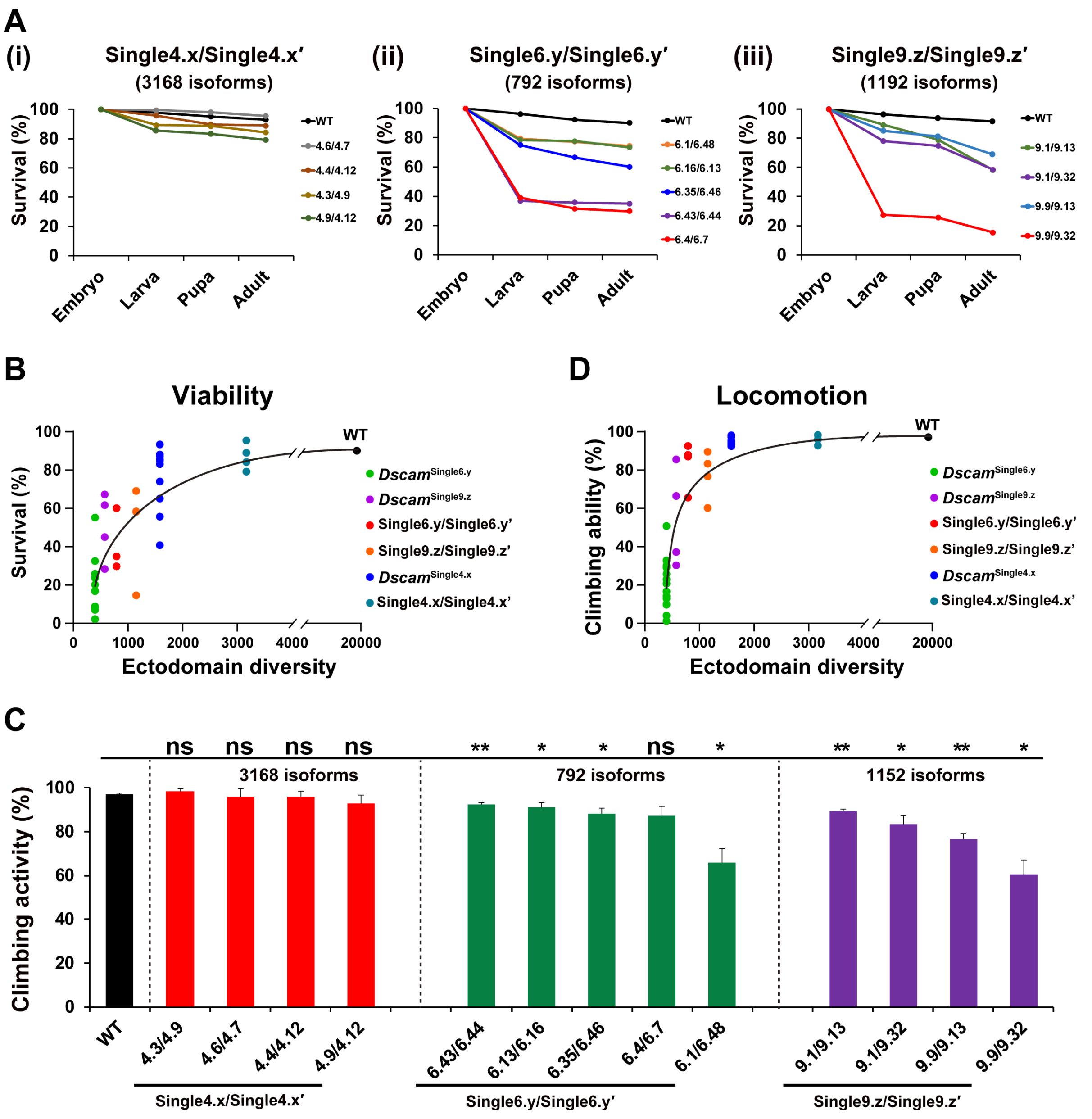


**Figure S2. The diversity of Dscam1 is required for *Drosophila* development. (Related to Figure 2).**

**(A)** Survival rates of wild-type and transheterozygous animals during development.

**(B)** The *Dscam1* isoform diversity is largely correlated with fly viability.

**(C)** Reducing Dscam1 diversity affects fly climbing ability. n.s., not significant; *P<0.05; **P<0.01; ***P<0.001. (Student's t-test, two-tailed).

**(D)** The *Dscam1* isoform diversity is largely correlated with fly climbing ability. However, these mutants with the same degree of diversity exhibited remarkable phenotypic differences, suggesting a cluster- and exon variant-specific manner.

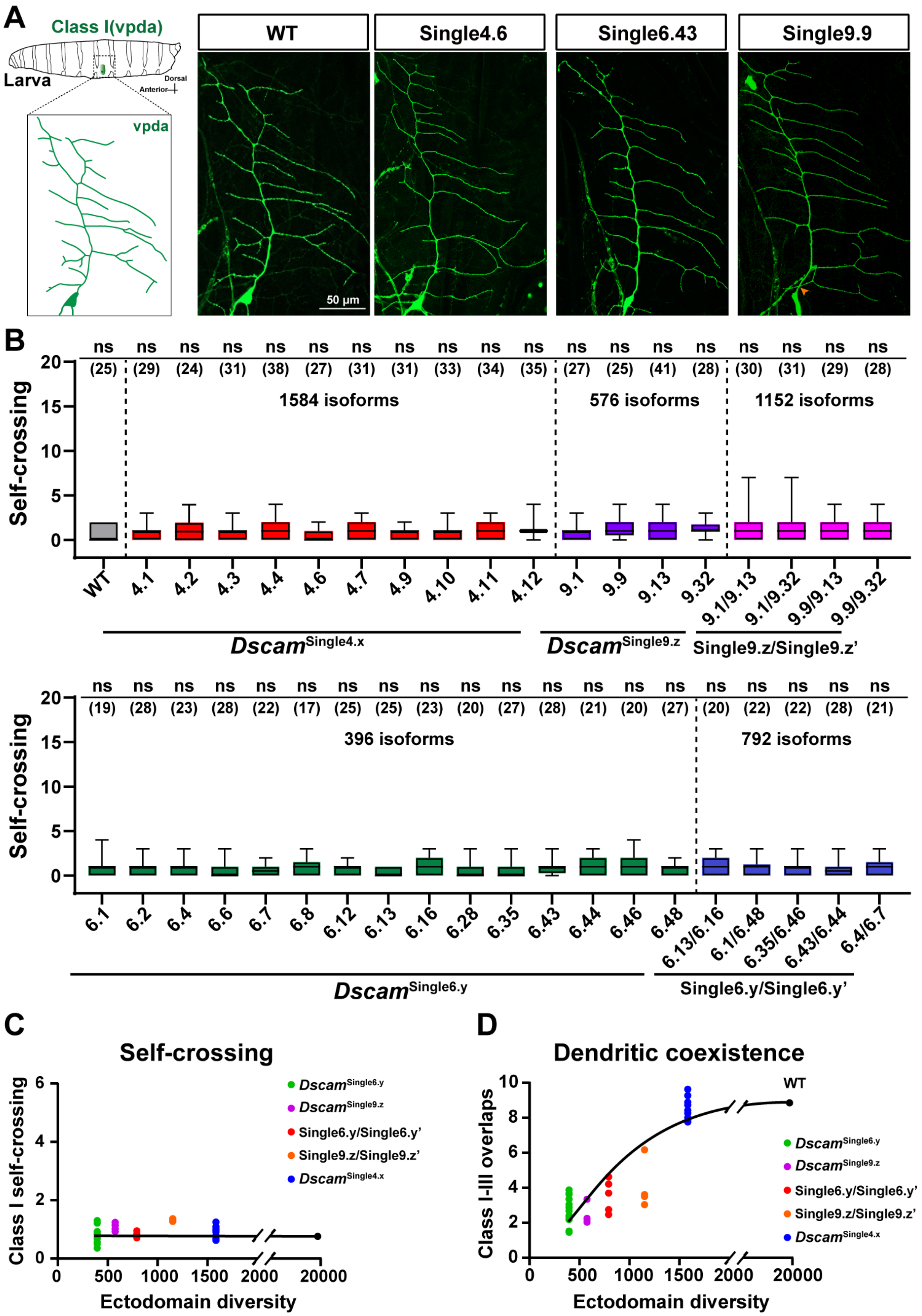


**Figure S3. Dscam1 diversity is dispensable for dendritic repulsion of class I da neurons. (Related to Figure 3).**

**(A)** Schematic diagram of class I neurons (vpda)**.** Representative images of dendrites of class I (vpda) neurons of different mutant flies. Scale bar, 50 µm.
**(B)** Self-repulsion was normal in all mutant flies similar to the wild-type control. Numbers in parentheses refer to the investigated neurons of each genotype. Data are expressed as mean ± SD. n.s., not significant (Student's t-test, two-tailed).
**(C)** Self-repulsion was normal independently of Dscam1 diversity.
**(D)** The dendritic coexistence between dendritic arborization neurons largely correlated with Dscam1 isoform diversity. However, these mutants with the same degree of diversity exhibited remarkable differences, suggesting a cluster- and exon variant-specific manner.

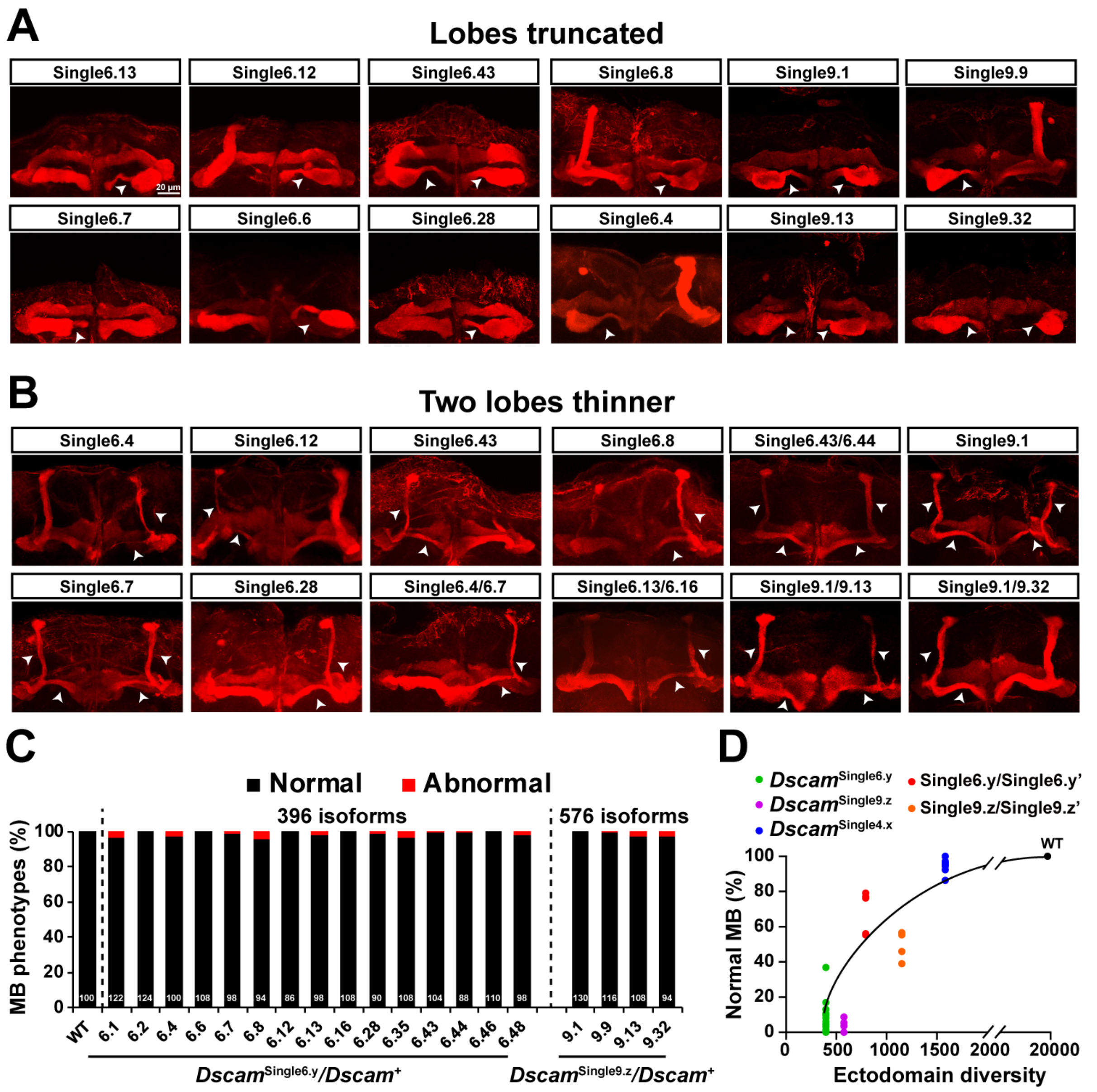


**Figure S4. Mushroom body phenotypes for *Dscam*$^{\mathrm{Single6.y}}$ and *Dscam*$^{\mathrm{Single9.z}}$ mutants. (Related to Figure 4).**

**(A)** The lobe truncation of the mushroom body in *Dscam*$^{\mathrm{Single6.y}}$ and *Dscam*$^{\mathrm{Single9.z}}$ mutants.

**(B)** The thinning of two lobes of the mushroom body in *Dscam*$^{\mathrm{Single6.y}}$ and *Dscam*$^{\mathrm{Single9.z}}$ mutants. Mushroom body lobe morphology in WT and *Dscam1* mutants were visualized with monoclonal antibody 1D4 (anti-Fas II, red). The white arrowheads indicate the defective lobes.

**(C)** MB phenotypes for *Dscam*$^{\mathrm{Single6.y}}$*/Dscam*$^{+}$ and *Dscam*$^{\mathrm{Single9.z}}$*/Dscam*$^{+}$ heterozygous mutants, which were indistinguishable from those of the wild-type control. Numbers in parentheses refer to the analyzed mushroom body neurons of each genotype.

**(D)** The MB phenotypes were largely correlated with Dscam1 isoform diversity.

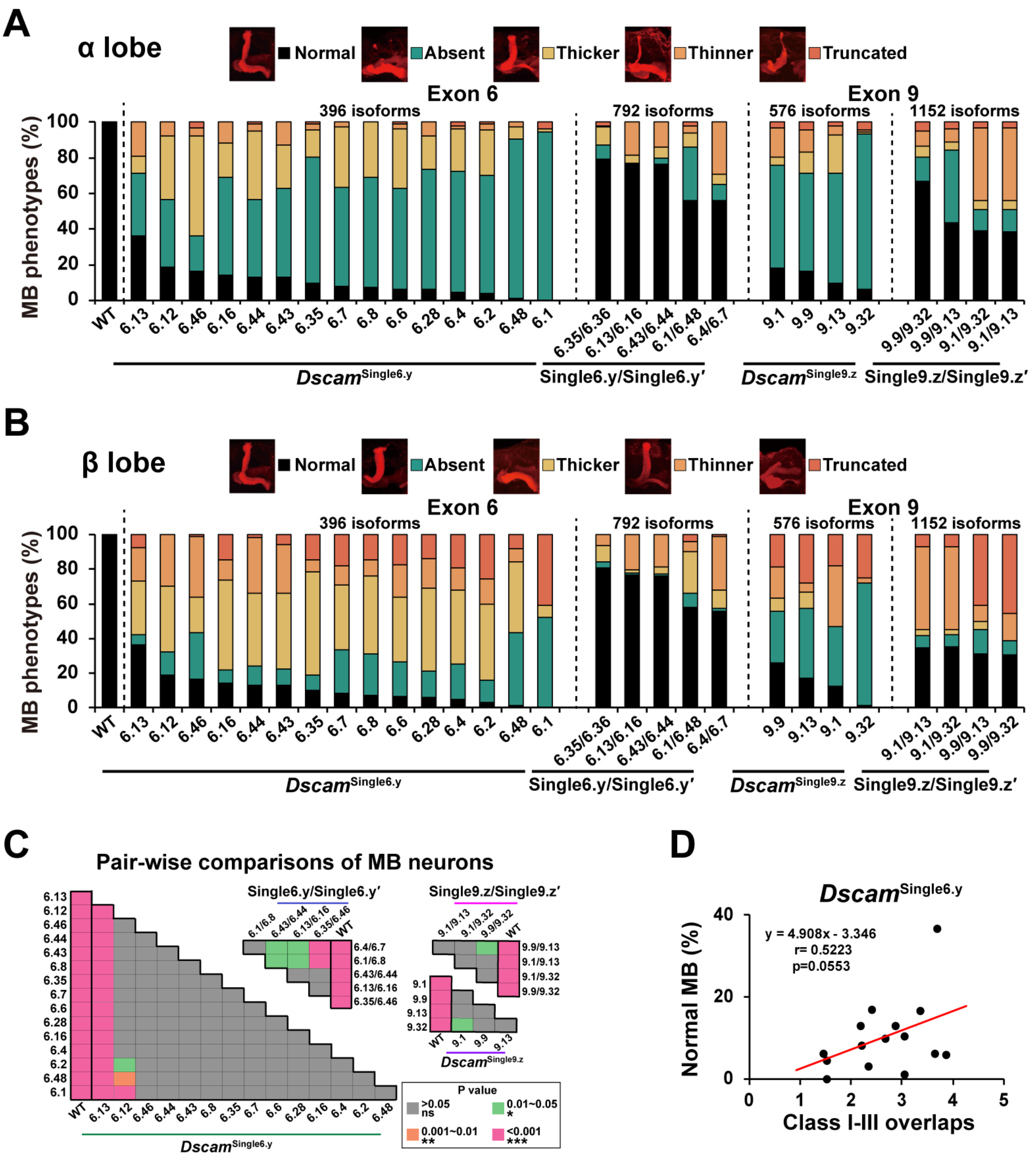


**Figure S5. Quantification of the MB defects in *Dscam*$^{\mathrm{Single6.y}}$ and *Dscam*$^{\mathrm{Single9.z}}$ mutants. (Related to Figures 4).**

**(A, B)** The defects of α (A) and β (B) lobes of MBs were analyzed. Although the α and β lobes of MBs did not exhibit obvious differences in overall defects, the spectra of individual defective phenotypes varied considerably. Our data showed lobe truncation mainly occurred in the β lobes of MBs.

**(C)** Pair-wise comparisons (one-way ANOVA with Tukey's test) of MB phenotypes among *Dscam*$^{\mathrm{Single6.y}}$ or *Dscam*$^{\mathrm{Single9.z}}$ mutants. These data indicate that mutants with the same degree of diversity exhibited phenotypic defect differences in MB phenotypes.

**(D)** MB phenotypes of individual *Dscam*$^{\mathrm{Singele6.y}}$ mutants showed no or marginally significant correlation with class I and III dendrite overlaps. This suggests that Dscam1 isoforms act on MB phenotypes via a pathway distinct from class I and III dendrite repulsion.

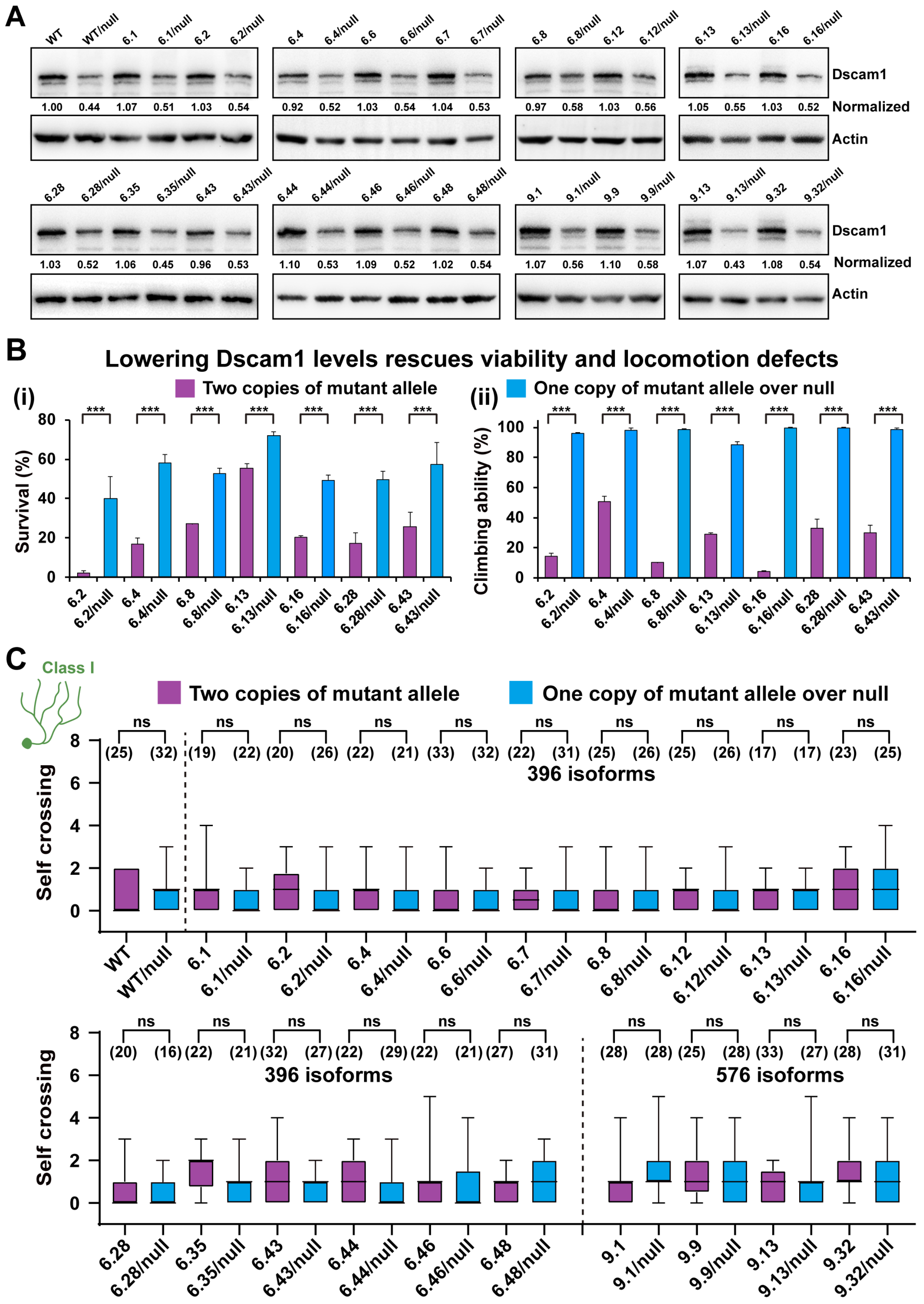


**Figure S6. Reducing Dscam1 expression level partially rescued the reduced fly viability and climbing ability in *Dscam*$^{\mathrm{Single6.y}}$ and *Dscam*$^{\mathrm{Single9.z}}$. (Related to Figure 7).**

**(A)** *Dscam*$^{Single6.y}$/*Dscam*$^{null}$ and *Dscam*$^{Single9.z}$/*Dscam*$^{null}$ flies reduced Dscam1 expression level by approximately 50% to wild-types. We used semi-quantitative Western blot analysis to compare Dscam1 protein levels between different genotypes. Dscam1 levels were normalized to β-actin levels, and the expression levels were then compared to the value of wild type, which was set to 1.

**(B)** Reducing Dscam1 expression level partially rescued the diminished fly viability (panel i) and climbing ability (panel ii) in homozygous mutants. Data are expressed as mean ± SD. *P < 0.05; **P < 0.01; ***P < 0.001; n.s., not significant (Student's t-test, two-tailed).

**(C)** Dendrites of class I neurons seldom exhibited self-crossing in *Dscam*$^{Single6.y}$/*Dscam*$^{null}$ and *Dscam*$^{Single9.z}$/*Dscam*$^{null}$ mutants similar to *Dscam*$^{Single6.y}$ and *Dscam*$^{Single9.z}$ mutants.

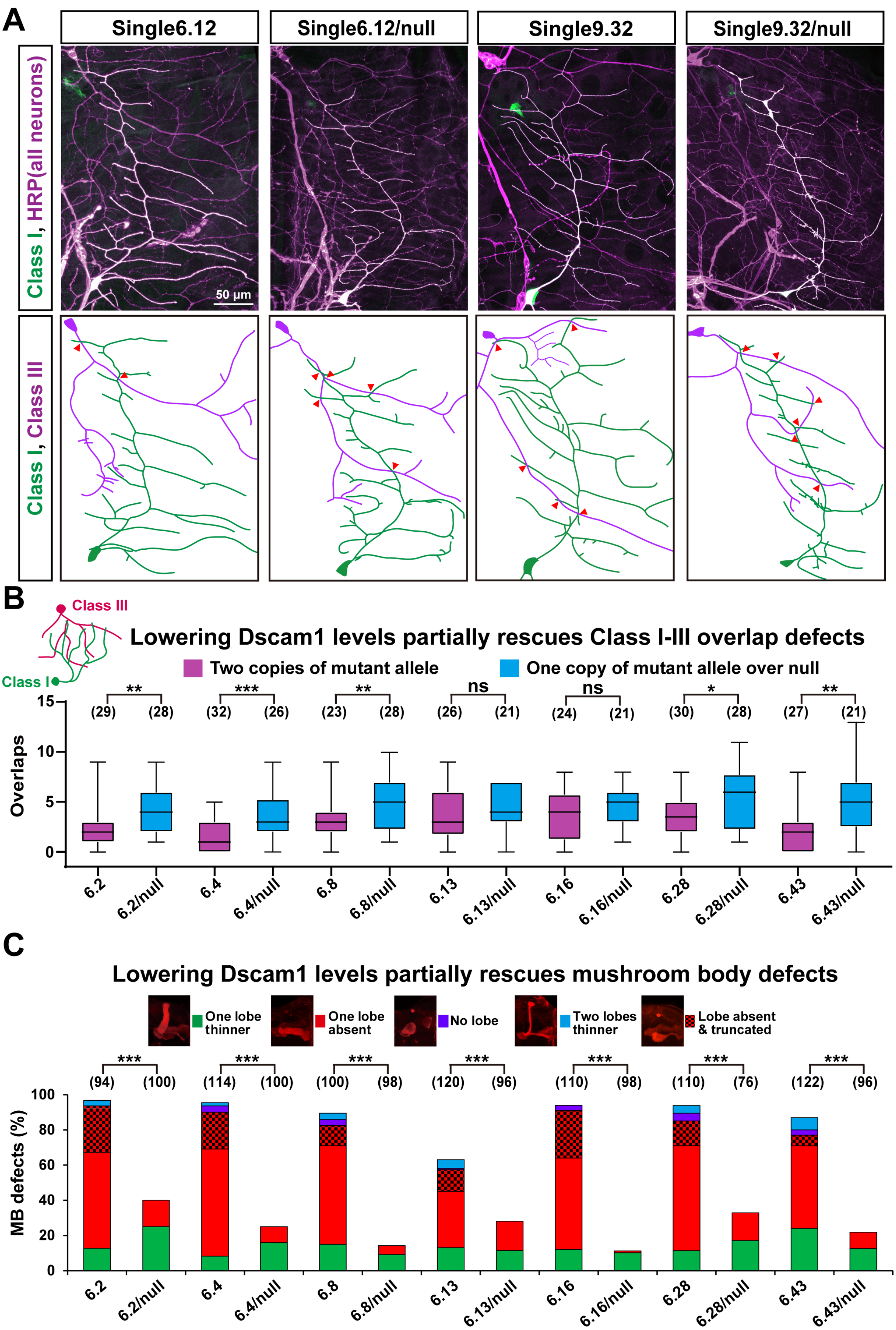


**Figure S7. Reducing the Dscam1 expression level partially rescues the neuronal defects. (Related to Figure 7).**

**(A)** Representative images of dendrites of dendritic arborization neurons in *Dscam1* mutants. Red arrowheads indicate the overlaps between class I and class III dendrites. Scale bar, 50 μm.

**(B)** Quantification of the overlaps between class I and class III dendrites in $Dscam^{Single6.y}/Dscam^{null}$ and $Dscam^{Single6.y}$ mutants. Numbers in parentheses represent the analyzed class I and class III neurons of each genotype. Data are expressed as mean ± SD. *P < 0.05; **P < 0.01; ***P < 0.001; n.s., not significant (Student's t-test, two-tailed).

**(C)** Quantification of the MB defects in $Dscam^{Single6.y}/Dscam^{null}$ and $Dscam^{Single6.y}$ mutants. Reducing Dscam1 expression level rescued mushroom body defects caused by reduced diversity in homozygous mutants. The numbers in the parenthesis represent the number investigated.
